# Temporal control of axonal floor plate crossing through a combination of incoherent feedforward and feedback loops of gene regulatory network regulating Robo3 expression

**DOI:** 10.1101/2025.10.26.684599

**Authors:** Reut Sudakevitz-Merzbach, Madhuri Majumder, Gerard Elberg, Marah Bakhtan, Dina Rekler, Sophie Khazanov, Benjamin J Wheaton, Nissim Ben-Arie, Gilgi Friedlander, Chaya Kalcheim, Sara Ivy Wilson, Alexander Jaworski, Miri Adler, Avihu Klar

**Author notes:** Correspnding authors: Miri Alder and Avihu Klar.

## Abstract

Spinal commissural neurons play a fundamental role in motor control and sensory perception. Robo3, a receptor expressed on pre-crossing commissural axons, is required for midline crossing. Its downregulation in post-crossing axons is essential for forming synapses with contralateral CNS targets. We demonstrate that, at the transcriptional level, the dynamic expression of *Robo3* in pre-crossing neurons is controlled by incoherent feed-forward loops (iFFLs) and negative feedback loops (NFLs) that precisely regulate its transience and thereby determine midline crossing. Lhx2 and Lhx9 activate *Robo3* while also inducing its repressor, Barhl2. Additionally, negative feedback loops fine-tune the transience of *Robo3* expression by adjusting the relative strength of the activation and repression modules. Our genomic analysis reveals that this regulatory circuitry converges on an enhancer element of the Robo3 gene. These findings imply that diverse iFFLs, together with NFLs, are essential for regulating the extent of midline crossing across different spinal commissural subtypes.

## Introduction

The formation of neuronal circuits requires the accurate navigation of axons to their proper targets. The interaction of axons with guidance cues depends on the temporal remodeling of axonal responsiveness. This is best exemplified by the midline crossing of commissural axons in the CNS (Chedotal, 2019; Kolodkin and Tessier-Lavigne, 2010). Commissural axons extend in a similar environment before and after crossing the midline; however, they respond antagonistically to the same molecular or cellular cues. For example, commissural premotor interneurons avoid forming synapses with motor neurons on the ipsilateral side but do so with motor neurons on the contralateral side (Moran-Rivard et al.; Pierani et al., 2001). Remodeling axonal responsiveness can be achieved post-translationally (through receptor shedding, receptor presentation, or interaction with guidance cues), post-transcriptionally (via RNA splicing or regulation of translation), or transcriptionally (Chedotal, 2019; Kolodkin and Tessier-Lavigne, 2010). Much is known about the role of transcription factors (TFs) in regulating the decision to cross or not cross the midline (Herrera and Escalante, 2022). However, little is known about how the gene regulatory network (GRN) contributes to the dynamic adaptation to guidance cues before and after midline crossing, and how defined classes of neurons change their laterality choice during development.

Entering, crossing, and exiting the floor plate (FP) requires rapid modulation of axonal responsiveness to FP-derived guidance cues. Attraction of pre-crossing commissural axons to Netrin1 changes to non-responsiveness in post-crossing axons (Shirasaki et al., 1998). Conversely, non-responsiveness to Slits in pre-crossing axons changes to repulsion in post-crossing axons (Zou et al., 2000). Robo3 is a key component in the transition from attraction to repulsion at the FP. In pre-crossing commissural axons, Robo3 expression is required for entry into the floor plate (Sabatier et al., 2004). Current understanding suggests that Robo3 silences Robo1 and Robo2, the receptors for Slits, and in mammals, Robo3 also mediates attraction as part of the DCC/Netrin1 signaling pathway (Zelina et al., 2014). The downregulation of Robo3 expression in post-crossing axons exposes them to the effects of Slits and ensures extension on the contralateral side of the floor plate (Jaworski et al., 2010). Thus, understanding how Robo3 is dynamically regulated in different populations is pivotal for determining how developmental circuitry is established.

The expression of Robo3 in commissural versus ipsilateral (ipsi) neurons, as well as in pre- and post-crossing commissural axons, is regulated at multiple levels: transcriptional, mRNA splicing, translational, and post-translational (Chedotal, 2019; Chen et al., 2008; Kolodkin and Tessier-Lavigne, 2010). At the transcriptional level, *Robo3* expression is regulated in the cardinal commissural interneurons by various TFs. The basic helix-loop-helix (bHLH) TFs NHLH1 and NHLH2 activate *Robo3* expression in all commissural neurons (Masuda et al., 2024). Additionally, the Lim-homeodomain (Lim-HD) TFs Lhx2 and Lhx9 activate expression specifically in dI1 neurons (Wilson et al., 2008), while the homeobox TF Dbx1 activates expression in dorsal commissural midbrain neurons (Inamata and Shirasaki, 2014). Conversely, *Robo3* is negatively regulated in some ipsilateral interneurons (INs). For instance, in the dIL_B_ subpopulation of ipsi-projecting spinal INs, Zic2 downregulates *Robo3* expression (Escalante et al., 2013). *Robo3* mRNA is expressed in early post-mitotic commissural INs, specifically in cells adjacent to the progenitor neuron domain, the ventricular zone. In late post-mitotic commissural INs, which migrate laterally, *Robo3* mRNA expression is downregulated as the leading process approaches the floor plate, a phenomenon demonstrated in the hindbrain precerebellar nuclei (Marillat et al., 2004). This indicates that the transition from upregulation to downregulation of *Robo3* is dynamically and temporally regulated. However, the mechanisms controlling the transience of *Robo3* expression, as well as whether the timing of *Robo3* expression in pre-crossing commissural axons influences floor plate crossing, remain open questions.

The laterality choice of the cardinal interneurons (INs) is robust. The majority of axons derived from the dI2, dI4, dI5, dI6, V0, and V3 populations are commissural, while most dI3, V1, and V2 axons are ipsilateral (Alaynick et al., 2011; Lai et al., 2016; Tulloch et al., 2019). dI1 neurons exhibit a temporally dynamic laterality choice, transitioning from commissural to ipsilateral. Early post-mitotic dI1 neurons are commissural, while late post-mitotic dI1 neurons give rise to two cardinal subpopulations with different axonal laterality choices: commissural dI1_comm_ and ipsilateral dI1_ipsi_ (Wilson et al., 2008). Post-gestation, most dI1 neurons are dI1_ipsi_ (Pop et al., 2022; Yuengert et al., 2015). Loss-of-function experiments in mice demonstrated that the commissural phenotype of dI1_comm_ is mediated by the TFs Lhx2 and Lhx9, which activate *Robo3* expression (Wilson et al., 2008). In contrast, Barhl2, a transcriptional repressor, is required for the ipsilateral projection (Ding et al., 2012). Barhl1, which is co-expressed with Barhl2 in dI1 neurons, is not required for the laterality choice of dI1 axons, as shown by the normal axonal phenotype in the spinal cord of Barhl1-null mice (Li et al., 2002). The finding that Lhx2, Lhx9, and Barhl2 expression is not simply segregated into dI1_comm_ and dI1_ipsi_ populations is inconsistent with a simple activation/repression model, in which activators are expressed in dI1_comm_ and repressors in dI1_ipsi_. Instead, both activators and the repressor are expressed in dI1_comm_ and dI1_ipsi_ neurons (Ding et al., 2012; Lee et al., 1998; Wilson et al., 2008).

The idea that the same cell can co-express both an activator and a repressor simultaneously, allowing each to transiently influence the cell’s phenotype, can be explained by the cell exploiting transcriptional regulatory networks (GRNs) of transcription factors (TFs). In this way, GRNs such as incoherent feed-forward loops (iFFLs) and negative feedback loops (NFLs) can function as switches for bimodal outputs—in this case, the on/off expression of a target gene. In a type 1 iFFL, a TF_activator_ upregulates the expression of a target gene while also inducing the expression of a TF_repressor_, which in turn represses the same target gene. This results in activation followed by suppression of the target gene (Mangan and Alon, 2003; Mangan et al., 2006). The robustness of an iFFL depends on the expression levels of the TF_activator_ and TF_repressor_, as well as the binding affinities of these TFs to cis-regulatory elements within the target gene’s enhancer. iFFLs are recurrent GRN patterns, also known as network motifs, that are enriched across diverse biological systems (Milo et al., 2002). Recently, we introduced the concept of network hyper-motifs to describe gene regulatory networks composed of multiple interconnected network motifs (Adler and Medzhitov, 2022). By combining elements such as iFFLs with feedback interactions, these hyper-motifs can enhance or fine-tune the functional capabilities of the individual motifs.

We hypothesize that Lhx2/9 and Barhl2 are part of a combined incoherent feed-forward loop (iFFL) and negative feedback loop (NFL) regulatory motif that governs the dynamic expression of Robo3 and, consequently, the axonal fate of dI1 neurons from dI1_comm_ to dI1_ipsi_. Using gain- and loss-of-function paradigms, we demonstrated that Lhx2/9 activate the expression of both Robo3 and Barhl2, while Barhl2 represses *Robo3* expression. The repression of *Robo3* is reinforced by additional regulatory modules: a negative feedback loop in which Barhl2 inhibits Lhx2/9, and a positive regulation of Barhl2 by Barhl1, with reciprocal reinforcement. We provide evidence that this iFFL converges on an evolutionarily conserved enhancer element of *Robo3*.

## Results

To track the temporal pattern of dI1 axonal projections, EGFP was stably expressed in dI1 neurons using a dI1-specific enhancer (edI1) and the Cre/LoxP system (Avraham et al., 2009; Hadas et al., 2014). Electroporation was performed in chicken embryos at HH Stage 18, and axonal choice between ipsilateral and contralateral projections was scored from E4 to E8. At E4, all axons projected toward the floor plate and crossed it contralaterally (Fig. 1A). From E4.5 onward, a gradual shift from contralateral to ipsilateral projections was observed (Fig. 1A–E).

**Fig. 1:**
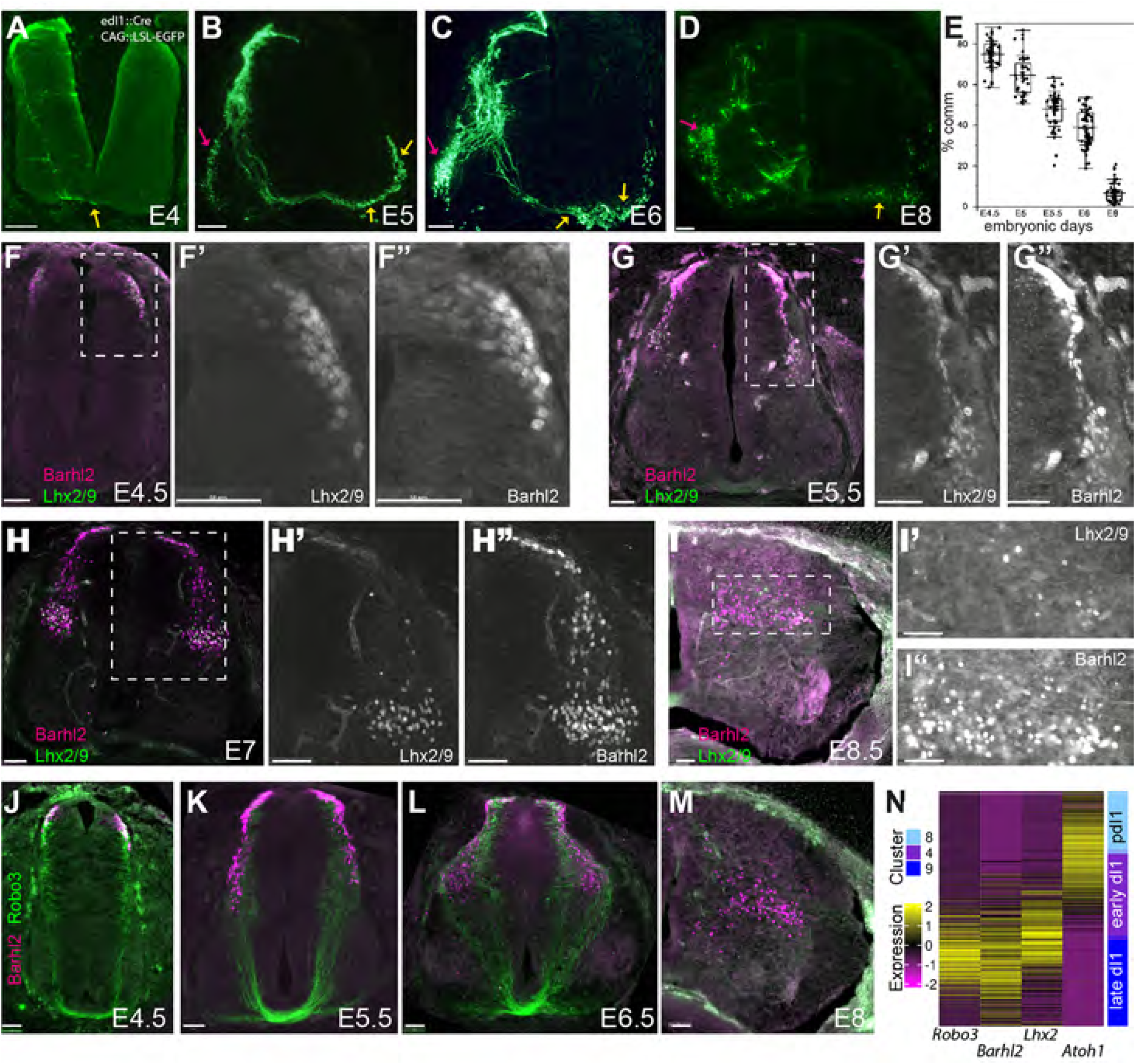
Axonal laterality choice and dynamic of transcription factors expression in dI1. A-D. Transvers sections of chick embryo spinal cord. Axonal projection of dI1 axons at E4 (A), E5 (B), E6 (C) and E8 (D) are shown. Yellow arrows - commissural axons, magenta arrows – ipsilateral axons. E. Box plot chart of the % of commissural axons. For each developmental stage (E4.5-E8), 10-15 cross sections from three embryos were analyzed (See Supplementary statistics). F-I. Transvers sections of chick embryo spinal cord. Expression patterns of Lhx2/9 and Barhl2 proteins from E4.5 to E8 are shown. J-M. Expression pattern of Robo3 and Barhl2 proteins from E4.5 to E7. N. Expression profiles of *Atoh1*, *Lhx2*, *Barhl2* and *Robo3* along the dI1 trajectory in the E4 Quail spinal cord (Rekler et al., 2024). Neurons are ordered by the pseudotime from up to down: cluster 8 – pdI1 progenitors (pdI1); cluster 4 – early differentiated dI1 (early dI1); cluster 9 – fully differentiated dI1 (late dI1). The heatmap displays log 2 transformed normalized counts, standardized per gene to have zero mean and unit standard deviation. Scale Bars = 50μm

The fate and axonal projections of dI1 neurons are characterized by the expression of the transcription factors (TFs) Atoh1, Barhl1, Barhl2, Lhx2, and Lhx9 (Lai et al., 2016). Atoh1, a bHLH transcription factor, is expressed in dI1 progenitors (pdI1) and activates the expression of post-mitotic TFs: the Lim-HD TFs Lhx2 and Lhx9, and the homeodomain TFs Barhl1 and Barhl2 (Herrera and Escalante, 2022). The co-expression of Lhx2/9 and Robo3 is consistent with Robo3 being positively regulated by Lhx2/9 (Fig. 1F–M). Barhl2 expression is evident as early as E4 - the commissural-only stage, and continues at E4–E6, when both commissural and ipsilateral neurons are present, and at E7/8, the ipsilateral stage (Fig. 1F–M). Thus, Barhl2 expression is sustained throughout development, including during the transition from commissural to ipsilateral axonal projection.

To test whether the commissural-to-ipsilateral temporal bias is also reflected at the level of mRNA expression of the TFs involved and *Robo3*, we analyzed single-cell RNA-seq data from the spinal cords of quail E4 (HH stage 23–24, comparable to chick E4–4.5) (Rekler et al., 2025; Rekler et al., 2024) and mouse E11.5 (Delile et al., 2019) (Fig. S1A). Pseudotime heatmap analysis was used to infer gene expression dynamics along the dI1 trajectory. Avian dI1 interneurons are represented in three discrete clusters, each characterized by specific gene expression signatures (Rekler et al., 2025) (Fig. S1A,B). In both mouse and quail, *Atoh1* is upregulated in progenitor dI1 (pdI1) and downregulated in early differentiated dI1. *Lhx2* and *Barhl2* are simultaneously upregulated in early differentiated dI1. *Lhx2* is downregulated in late differentiated dI1, while *Barhl2* expression is maintained (Fig. 1N, Fig. S1D,E). The transient expression profile of *Robo3* is similar to that of Lhx2, though with a slight temporal delay (Fig. 1N, S1A). The negative correlation between *Barhl2* and *Robo3* at later stages (Fig. 1M) supports the role of Barhl2 as a negative regulator of Robo3. However, the co-expression of *Lhx2/9* and *Barhl2* at both the commissural-only stage (Fig. 1F) and the commissural-to-ipsilateral transition stage (Fig. 1G) challenges a simple direct activation/repression model. We propose that Lhx2/9 and Barhl2 regulate Robo3 expression through an incoherent feed-forward loop (iFFL), in which Lhx2/9 activate both Robo3 and Barhl2, while Barhl2 represses Robo3 (Fig. 2A). This temporal GRN can explain the early *Robo3*^+^ and later *Robo3*^-^ expression, and consequently the shift from commissural to ipsilateral projection.

**Figure 2:**
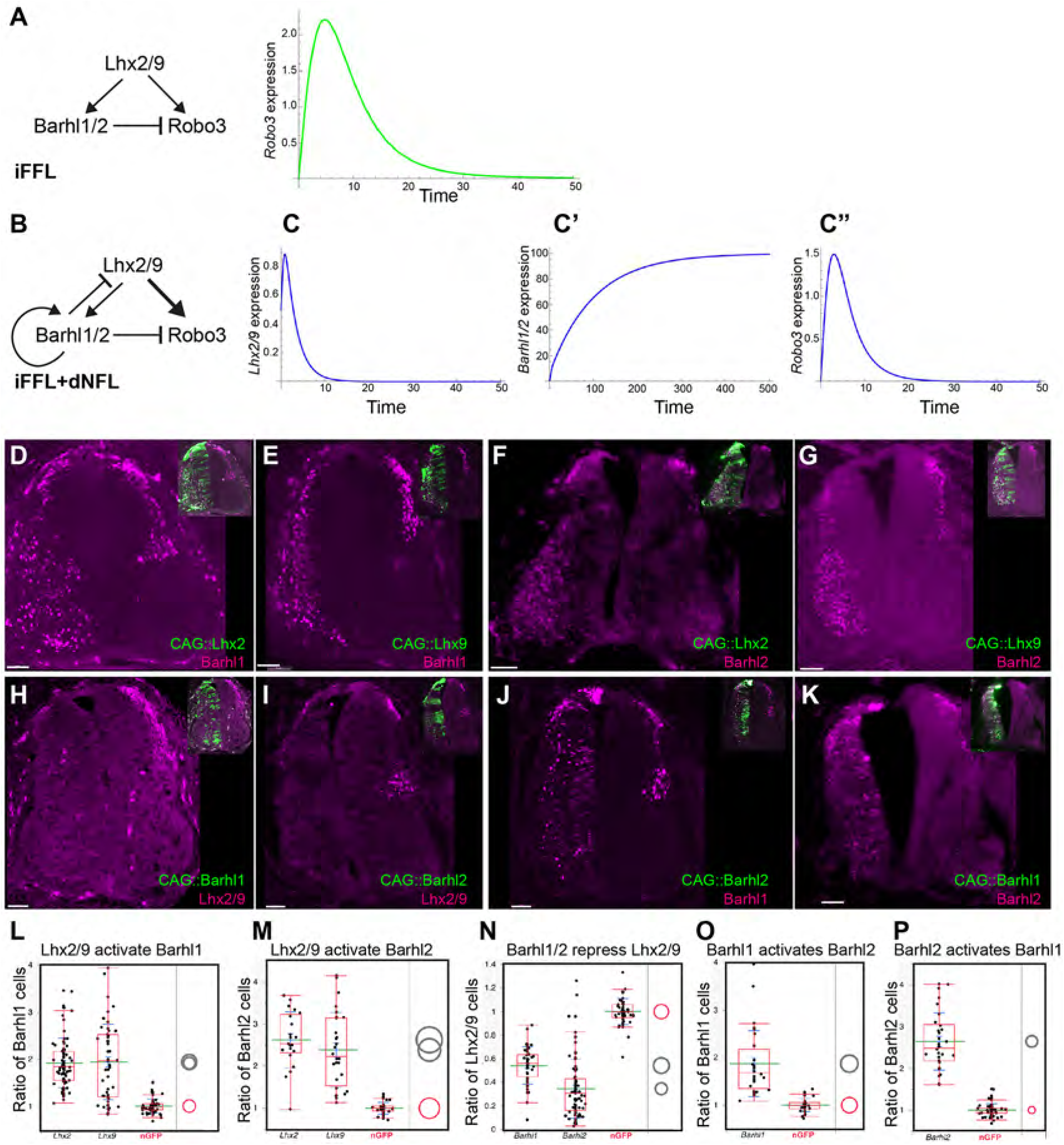
Gene regulatory network of Lhx2/9, Barhl1/2 and *Robo3* in the spinal cord. A. Time-course simulations of *Robo3* expression via type 1 incoherent feedforward loop (iFFL) where Lhx2/9 activates Barhl1/2, which represses *Robo3*. This configuration yields a transient Robo3 response. B. Schematic of the core gene regulatory network motif in which Lhx2/9 activates both *Robo3* and its repressor *Barhl1/2*. Barhl1/2 negatively regulates *Robo3* and provides inhibitory feedback to *Lhx2/9* while also engaging in positive autoregulation (iFFL with double feedback loops – iFFL+dNFL). C. Time-course simulations of *Lhx2/9* (C), *Barhl1/2* (C’), and *Robo3* (C”) expression levels for the iFFL+dNFL network. The system exhibits a transient activation of Robo3, followed by stabilization of Barhl1/2 and reduction of Lhx2/9. For all the experiments in D-K, a transcription factor was electroprated into the hemi-tube at HH st. 18, and the embryo were analyzed at E5. The TFs are cloned upstream to IRES-nEGFP cassette in pCAG vector. The number of the target TF in the electroporated and control sides were scored. The ratio between the two sides was calculated. In all the images, the pattern of the ectopically expressed TF (EGFP+) is shown in the insets. D-K. Transverse sections of E5.5 chick spinal cords. Lhx2 (E) and Lhx9 (F) activate Barhl1 (D,E). Lhx2 (G) and Lhx9 (H) activate Barhl2 (F,G). Barhl1 (I) and Barhl2 (J) inhibit Lhx2/9 (H,I). Barhl1 and Barhl2 activate each other (J,K). L. Quantification of the ratio of Barhl2 cells between the Lhx2 and Lhx9-electroporated and the control sides. The mean ratios are the following: nGFP – 1+/-0.14; Lhx2 – 1.92+/-0.52; Lhx9 – 1.94+/-0.8. M. Quantification of the ratio of Barhl1 cells between the Lhx2 and Lhx9-electroporated and the control sides. The mean ratios are the following: nGFP – 0.99+/-0.13; Lhx2 – 2.62+/-0.66; Lhx9 – 2.38+/-0.88. N. Quantification of the ratio of Lhx2/9+ cells between the Barhl1/2-electroporated and the control sides. The mean ratios are the following: nGFP – 1+/-0.11; Barhl1 – 0.54+/-0.15; Barhl2 – 0.34+/-0.25. O. Quantification of the ratio of Barhl2 cells between the Barhl1-electroporated and the control sides. The mean ratios are the following: nGFP – 0.99+/-0.13; Barhl1 – 1.87+/-0.69. P. Quantification of the ratio of Barhl1 cells between the Barhl2-electroporated and the control sides. The mean ratios are the following: nGFP – 1+/-0.14; Barhl2 – 2.64+/-0.68. For each electroporation, cross section from three embryos were analyzed, 10-15 sections from each embryo. Comparing the electroporated/control-nonelectroporates ratios across groups using Dunnett’s method (which takes into account multiple comparisons) shows significant differences between the electroporated and control-nonelectroporates in all experiments. The circle charts show the significance. No overlapping between the experimental electroporated side (gray circles) and control-nonelectroporates (red circle) is indicative of *P* < 0.05 (see Supplementary Statistics). Scale Bars = 50μm

Incoherent feed-forward loops (iFFLs) have been widely studied for their dynamic properties across biological systems (Lee et al., 2002; Mangan and Alon, 2003; Milo et al., 2002). They play important roles in developmental programs and have been identified in transcriptional networks of human embryonic and hematopoietic stem cells, as well as downstream of the Notch signaling pathway (Boyer et al., 2005; Swiers et al., 2006). Key features of iFFLs include accelerated response times of the target gene (Mangan and Alon; Mangan et al., 2006) and fold-change detection, in which the system responds to relative rather than absolute changes in the input signal (Adler et al., 2018; Goentoro et al., 2009). Additionally, iFFLs can generate transient, pulse-like expression of the target gene, in this case, *Robo3*. Our data revealed that mathematical modeling of an iFFL circuit involving Lhx2/9 and Barhl1/2 (inputs) and *Robo3* (output) recapitulates this behavior, producing the expected transient expression profile of *Robo3* (Fig. 2A).

### Lhx2/9 Positively Regulate Barhl1/2, While Barhl1/2 Repress Lhx2/9

We tested the validity of the iFFL GRN hypothesis by examining the influence of each transcription factor (TF) on the others. Following ectopic expression of Lhx2 or Lhx9 in the chick spinal cord, the expression of Barhl1 and Barhl2 proteins was scored on both the electroporated and non-electroporated sides. The ratio of Barhl1/2-expressing cells between the two hemispheres of the spinal cord was then calculated. The number of Barhl1- and Barhl2-expressing cells increased significantly following ectopic expression of Lhx2 or Lhx9 (Fig. 2D–G, L, M). To test whether Lhx2/9 are required for Barhl2 expression, we examined Barhl2 expression in the spinal cords of *Lhx2/9* double knockout mouse embryos compared with control heterozygote or single homozygote littermates at E11.5 brachial spinal cord, a stage when Barhl2 is clearly expressed in Lhx2/9^+^ neurons (Wilson et al., 2008). Barhl2 expression was detected in both double knockout (n = 3 embryos) and littermate control (n = 9 embryos) embryos, with Barhl2^+^ cells present in similar numbers regardless of genotype (Fig. S2A–D). Interestingly, a statistically significant trend toward lower Barhl2 expression levels was observed in double knockouts compared with controls, supporting the idea that Lhx2 and Lhx9 contribute to Barhl2 regulation (Fig. S2E,F). Thus, at E11.5, loss of Lhx2/9 function does not result in complete loss of Barhl2 expression. We conclude that Barhl2 expression is regulated via a coherent feed-forward loop: Atoh1 activates both Lhx2/9 and Barhl2 (Reig et al., 2007; Saba et al., 2005). Lhx2/9 also activate Barhl2 expression, which is further maintained by Barhl1 (see below). The early onset of expression by Atoh1, combined with autoregulation by Barhl proteins, is sufficient to sustain Barhl2 expression, albeit at reduced levels, in the absence of Lhx2/9.

Ectopic expression of either Barhl1 or Barhl2 reduced the number of Lhx2/9-expressing cells (Fig. 2H, I, N), supporting a role for Barhl1/2 in suppressing *Lhx2/9*. Barhl1/2 also positively regulate their own expression, as ectopic expression of Barhl1 or Barhl2 induced expression of the paralogous gene (Fig. 2J, K, O, P). The positive autoregulation of Barhl TFs is consistent with the reduced Barhl1 levels observed in the spinal cords of Barhl2-null mice (Ding et al., 2012). These gain-of-function experiments support the activation module of the iFFL proposed in this study. Furthermore, our analysis revealed two additional feedback modules integrated into the iFFL design: (1) negative feedback, in which Barhl1/2 repress Lhx2/9, and (2) positive autoregulation of Barhl1/2. We therefore suggest that in dI1 neurons, Robo3 is regulated by a combined iFFL and NFL design (Fig. 2B).

### Barhl2 Inhibits Midline Crossing by Repressing Robo3 Expression

To test whether the transition from commissural to ipsilateral axonal projection is mediated by Barhl2, we ectopically expressed Barhl2 in dI1 neurons. In control experiments, dI1 neurons expressing mCherry showed that 48 ± 18% of axons were commissural (Fig. 3A, D). Upon ectopic expression of Barhl2, only 10 ± 6% of axons were commissural (Fig. 3B, D). The commissural-to-ipsilateral transition induced by Barhl2 was rescued by co-expression of Robo3 in dI1 neurons, supporting the notion that Barhl2 suppresses *Robo3* expression. In fact, most dI1 axons (64 ± 17%) co-expressing Barhl2 and Robo3 crossed the midline (Fig. 3C, D). Thus, as predicted by the model, Barhl2 gain-of-function has the opposite effect of the commissural-only phenotype observed in Barhl2-null mice (Ding et al., 2012). Therefore, high levels of Barhl2 are sufficient to inhibit midline crossing.

**Figure. 3.**
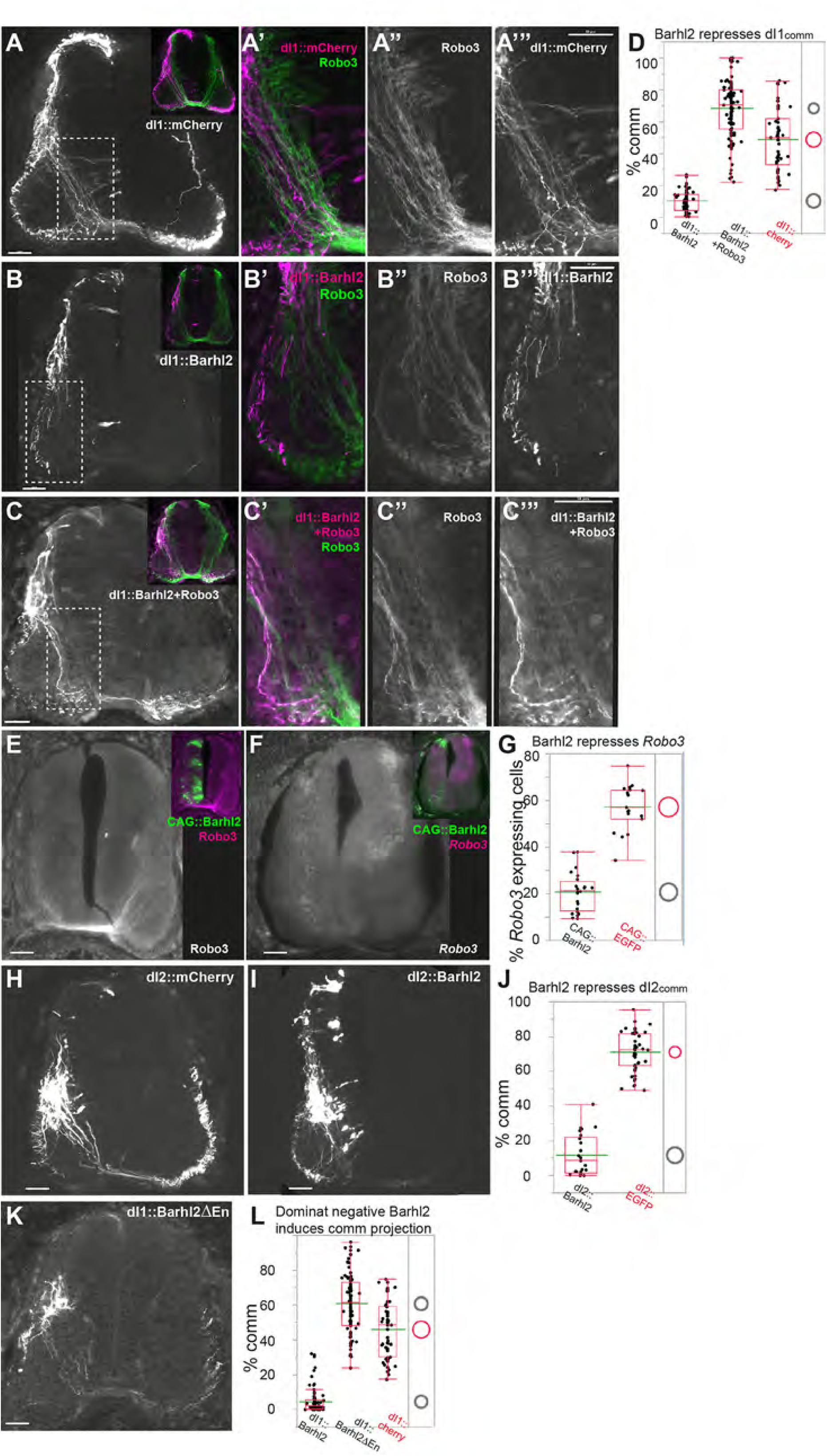
Barhl2 inhibits midline crossing via repression of Robo3. For interneurons subtype specific expression (A-C, I-,L), Barhl2 and its isoforms were cloned upstream to IRES-mCherry cassette in a Cre-conditional CAG::LSL plasmid. For targeting dI1 (A-C,L) or dI2 neurons (I,J), the Barhl2 plasmids were co-electroporated with dI1::Cre plasmid or dI2::Cre plasmid, respectively. For uniform ectopic expression (E-G), Barhl1 and Barhl2 under the control of CAG, were electroporated. For all the experiments, the plasmids were electroprated into the hemi-tube at HH st. 18, and the embryo were analyzed at E5.5. For each electroporation, 10-15 cross sections from three embryos were analyzed. A-C: Axonal projection following ectopic expression of Barhl2 in dI1. The dotted rectangles are enlarged in the three images at the right sides. A. dI1::mCherry - axons project ipsilaterally and contralaterally. B. dI1::Barhl2 - ipsilaterally bias. C. dI1::Barhl2+Robo3 - contralaterally bias. D. Quantification of the ratio of dI1 commissural axons from the total number of axons. Comparing the ratio of commissural axons between the control – dI1::mCherry, and the experimental dI1::Barhl2 and dI1::Barhl2+Robo3, using Dunnett’s method shows significant differences between the control in all experiments *P* < 0.05. (see Supplementary Statistics). E. Expression of Robo3 following Barhl2 ectopic expression. insets Barhl2+EGFP expressing cells. F. Expression of *Robo3* mRNA following Barhl2 ectopic expression. G. Quantification of the ratio of *Robo3* mRNA expression following ectopic expression of Barhl2. Comparing the ratio of Robo3 mRNA between the control – CAG::GFP, and the experimental CAG::Barhl2, using Dunnett’s method shows significant differences between the control and CAG::Barhl2 *P* < 0.05 (see Supplementary Statistics). H,I. Axonal projection following ectopic expression of Barhl2 in dI2. In control experiment, most of EGFP expressing dI2 axons project contralaterally (H). An ipsilaterally bias of dI2::Barhl2 axons (I). J. Quantification of the ratio of dI2 commissural axons following ectopic expression of Barhl2 from the total number of axons. Comparing the ratio of commissural axons between the control – dI2::mCherry, and the experimental dI2::Barhl2, using Dunnett’s method shows significant differences between the control and dI2::Barhl2 *P* < 0.05. (see Supplementary Statistics). K. A commissural projection bias following ectopic expression of a dominant negative isoforms that lacks the En recruitment domains - Barhl2ΔEn. L. Quantification of the ratio of dI1 commissural axons following ectopic expression of Barhl2’s isoform from the total number of axons. Comparing the ratio of commissural axons between the control – dI1::mCherry, and the experimental dI1::Barhl2 and dI1::Barhl2DEn, using Dunnett’s method shows significant differences between the control in all experiments *P* < 0.05. (see Supplementary Statistics). Scale Bars = 50μm

Inhibition of midline crossing can be achieved by repression of *Robo3* alone, making Robo3 central to this fate choice (Sabatier et al., 2004). We therefore examined Robo3 expression following ectopic expression of Barhl2 using in situ hybridization and immunostaining. *Robo3* mRNA and protein levels were downregulated on the electroporated side following Barhl2 expression (Fig. 3E–G). The repression of Lhx2/9 and Robo3 by Barhl2 does not exclude the possibility that in dI1 neurons, Lhx2/9 are the primary targets of Barhl2, and that repression of midline crossing is a secondary effect mediated by downregulation of Lhx2/9. Supporting this idea, the ipsilateral-bias phenotype induced by Barhl2 ectopic expression was rescued by co-expression of either Lhx2 or Lhx9 in dI1 neurons (Fig. S3). To test whether Barhl2 can repress midline crossing directly, we expressed Barhl2 in the commissural cardinal population dI2. We used the *Foxd3* enhancer, which drives expression in dI2 and V1 neurons (Avraham et al., 2009; Haimson et al., 2021). To increase expression specifically in dI2 neurons, electroporation was targeted to the dorsal spinal cord. In control experiments, 71.1 ± 11% of dI2::GFP axons were commissural. In contrast, in dI2::Barhl2 neurons, only 11.7 ± 11% of axons were commissural (Fig. 3H–J). dI2 neurons express Lhx1 and Lhx5, which could theoretically substitute for Lhx2/9 in dI2, but their expression was not affected by Barhl2 (Fig. S5).

To test these findings from another perspective, we used a loss-of-function paradigm by expressing a dominant-negative isoform of Barhl2 in dI1 neurons. Barhl2 is a Groucho-dependent transcriptional repressor that binds DNA via a homeodomain module and recruits Groucho through two Engrailed (En) recruitment domains (Goldstein et al., 2005; Sena et al., 2019). Mutation of the En domains preserves DNA binding but prevents Groucho recruitment, creating a dominant-negative allele (Sena et al., 2019). Ectopic expression of Barhl2ΔEn in dI1 neurons caused a shift in axonal projections from ipsilateral to commissural (Fig. 3K, L), similar to the phenotype observed in Barhl2-null mice. We conclude that Barhl2 inhibits midline crossing in dI1 neurons by repressing Robo3 and consolidates the ipsilateral projection by repressing Lhx2/9.

### Feedback-Enhanced iFFL Architecture Optimizes the Timing of Robo3 Expression

To better understand how the proposed regulatory architecture shapes *Robo3* expression dynamics, we developed a mathematical model of the circuit in which Lhx2/9 activate both *Robo3* and its repressor *Barhl1/2*. Barhl2, in turn, directly represses *Robo3*, inhibits *Lhx2/9* through a negative feedback loop, and reinforces its own expression via a positive autoregulation loop. This results in a combined iFFL with double feedback loops (iFFL+dNFL) (Fig. 2B). Simulations of the iFFL+dNFL circuit reproduced the hallmark dynamic signature of an iFFL: an initial activation of Robo3 followed by a rapid decline, yielding a transient, pulse-like expression profile. In the model, Lhx2/9 expression rises quickly, driving early *Robo3* induction, while Barhl1/2 accumulate more slowly. Once Barhl2 reaches sufficient levels, it represses *Robo3* expression, thereby terminating the pulse (Fig. 2B, C). This behavior matches our experimental observations that Robo3 expression is transient during development, with high levels at early stages and downregulation at later stages.

Although a simple iFFL design can also generate a transient *Robo3* response (Fig. 2A, Fig. 4A), our simulations comparing the observed circuit to three alternative designs, each lacking one or both feedback interactions, showed that the Negative Feedback from Barhl1/2 to Lhx2/9 (iFFL+_BL_NFL) and the Positive Autoregulation of Barhl1/2 (iFFL+_BB_PA) each make distinct and complementary contributions to shaping the *Robo3* pulse. The negative feedback provides tighter control over pulse duration by shusng down the activator Lhx2/9 early, preventing sustained *Robo3* expression. In contrast, the positive autoregulation of Barhl1/2 provides tighter control over the amplitude of the *Robo3* response by accelerating the accumulation of Barhl1/2, the repressor of *Robo3*, thereby ensuring rapid repression once the pulse is initiated. Incorporating both feedback loops (iFFL+dNFL) produced the most precisely timed and amplitude-controlled *Robo3* pulse, sharpening its onset, accelerating its termination, and reducing variability in both duration and magnitude compared to the simpler iFFL (Fig. 4A–C, Fig. S6). Together, the modeling results indicate that the combined iFFL, negative feedback, and positive autoregulation loops are optimal for generating the sharply timed window of *Robo3* expression required for proper commissural-to-ipsilateral axonal switching.

**Figure 4:**
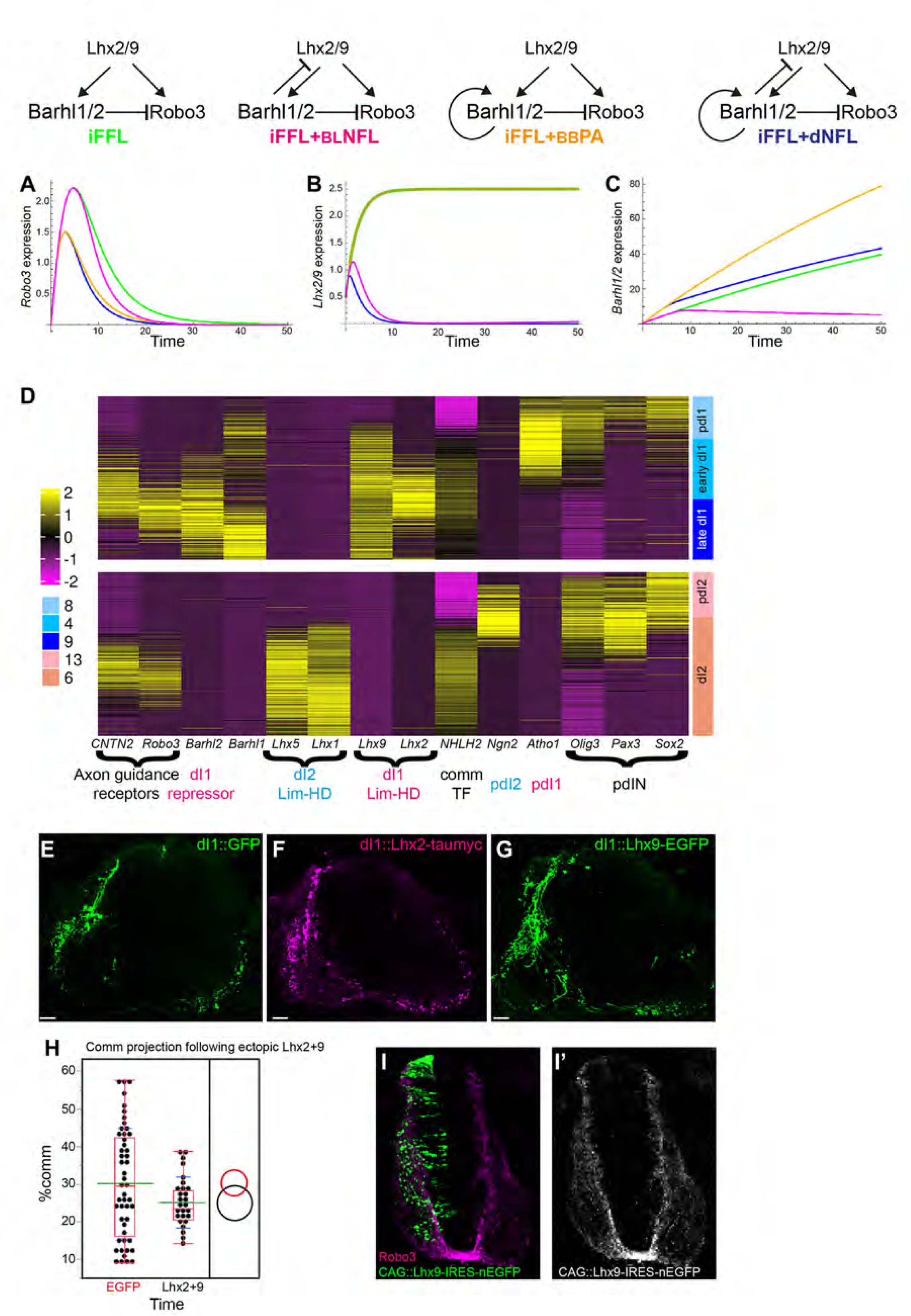
Network architecture and dynamic behavior of Lhx2/9–Barhl1/2–Robo3 regulatory motifs. A-C. Comparative dynamics of *Robo3*, *Lhx2/9*, and *Barhl1/2* across four distinct network topologies (color-coded – see below), each incorporating different combinations of feedback and regulatory interactions. The corresponding schematic diagrams are shown on the right. Plots display the temporal expression profiles of Robo3 (A), Lhx2/9 (B), and Barhl1/2 (C) for each topology: simple i1FFL iFFL (green). iFFL plus Barhl1/2 negative feedback to Lhx2/9 - iFFL+BLNFL (magenta). i1FFL plus Barhl1/2 positive autoregulation - iFFL+BBPA (orange). i1FFL plus Barhl1/2 self-activation and negative feedback to Lhx2/9 - iFFL+dNFL (blue). D. Expression profiles of selected genes along the dI1 and dI2 trajectories of E4 Quail spinal cord. For the dI1 trajectory, cells were randomly subsampled to have the same number of cells in dI1 and dI2 trajectories (478 cells). For each trajectory, the cells are ordered by the pseudotime, and cluster identities are indicated in the top annotation (Fig. S1B,C). The heatmap displays log 2 transformed normalized counts, standardized per gene to have zero mean and unit standard deviation. E-H. Transvers sections of embryonic chick spinal cord following ectopic expression of Lhx2 and Lhx9 in dI1. The following combinations of plasmids was used: dI1::Cre (E-G), CAG::LSL-EGFP (E), CAG::LSL-Lhx2-IRES-taumyc (F), CAG::LSL-Lhx9-IRES-EGFP (G). H. Quantification of extent of ipsilateral turning in dI1 following Lhx2+Lhx9 expression in dI1 at E6. Comparing the % of ipsilateral axonal projection between control-EGFP to the experimental Lhx2/9 across groups using Dunnett’s method (which takes into account multiple comparisons) shows no significant differences between the EGFP and the Lhx2/9 expressing dI1 neurons. (see Supplementary Statistics). H. Expression of Robo3 following etopic expression of Lhx9. Scale Bars = 50μm

The temporal expression of Robo3 is a feature shared by all commissural neurons (Delile et al., 2019; Marillat et al., 2004; Sabatier et al., 2004). Downregulation of Robo3 in post-crossing axons is required to prevent stalling at the floor plate and to enable axonal trajectories toward post–floor plate targets, through the transition of axonal responses from attraction to repulsion at the midline (Chen et al., 2008; Sakai and Kaprielian, 2012). Thus, a similar iFFL may control the transient temporal expression of *Robo3* in other commissural cardinal subtypes of the spinal cord. To test whether the transcription factor (TF) profiles that regulate cell fate and *Robo3* expression are similar in other commissural neurons, we compared their expression in dI1 and dI2 neurons (Fig. 4D, Fig. S1A–C). Trajectory and pseudotime heatmap analyses revealed that the earliest onset of expression in both dI1 and dI2 neurons involves the dorsal progenitor TFs *Pax3* and *Olig3*, along with the neuronal progenitor TF *Sox2*. Next, cell type– specific progenitor bHLH TFs are expressed: *Atoh1* in pdI1 and *Ngn2* in pdI2. Post-mitotic dI1 and dI2 neurons express the activators of *Robo3*—Lim-HD and bHLH TFs. *NHLH2*, a bHLH TF that regulates Robo3 expression in all commissural neurons (Masuda et al., 2024), is expressed in both dI1 and dI2 in a similar temporal pattern. Lim-HD TFs are expressed in early post-mitotic neurons: *Lhx2/9* in dI1 and *Lhx1/5* in dI2. The transient expression profiles of the pre-crossing receptors *Robo3* and *TAG1/CNTN2* are also similar in dI1 and dI2. Thus, the temporal expression patterns of *Robo3* and its regulators are similar in dI1 and dI2, suggesting that an iFFL, with an as-yet unidentified repressor, may control Robo3 expression in dI2 and other commissural neurons. The commissural dominance in dI2, as opposed to the ipsilateral dominance in dI1, may result from a variant iFFL—such as the basic iFFL or iFFL+_BL_NFL (Fig. 4A), that enables a more extended period of transient Robo3 expression, thereby facilitating midline crossing.

### Removal of the Negative Feedback Component from the GRN of Robo3 Does Not Alter Midline Crossing

Among the four possible iFFL designs, the two that provide the tightest control over pulse duration are iFFL+_BB_PA and iFFL+dNFL (Fig. 4A). It is predicted that removal of the negative feedback inhibition of Barhl2 over *Lhx2/9* would not change the amplitude of *Robo3* expression and, consequently, would not alter the extent of midline crossing. To test this hypothesis, Lhx2/9 were ectopically expressed in dI1 neurons under the control of the ubiquitously expressed CAG enhancer/promoter, which is not subject to Barhl2 regulation. At E6, 72 hours post-electroporation, no significant difference in commissural versus ipsilateral axons was observed between EGFP controls and ectopic Lhx2/9-expressing neurons (Fig. 4E–H). Interestingly, a transient bias toward ipsilateral projection was evident at E5, 48 hours after electroporation of Lhx2/9 (Fig. S4). This temporary wave of ipsilateral projections may reflect the robust transcriptional activation of endogenous *Barhl2* by ectopic Lhx2/9, which is subsequently balanced by cumulative Lhx2/9 levels. A similar ipsilateral bias was observed when Lhx2 or Lhx9 were ectopically expressed in all spinal neurons (Fernandez-Nogales et al., 2022).

The inability of ectopic Lhx2/9 expression to impose additional midline crossing is also reflected in the lack of Robo3 induction. As predicted by our mathematical modeling, the number of Robo3^+^ axons following Lhx9 ectopic expression was similar on the electroporated and control sides (Fig. 4I). Therefore, removal of the negative feedback module from the GRN does not disturb Robo3 expression or midline crossing.

### Lhx2/9 Activate Robo3 Expression by Direct Binding to Cis-Regulatory Elements Within the Robo3 Enhancer

To test whether the transcription factors (TFs) within the Robo3 iFFL exert their activity directly on the *Robo3* gene, we next aimed to identify the *Robo3* gene enhancer element. We cloned 5.5 kb (eR3-123 clone in Fig. 5A) of the chick *Robo3* upstream region into an mCherry reporter plasmid. Expression of the reporter gene in transgenic chick embryos yielded exclusive expression in commissural neurons (Fig. 5B, H). Using deletion constructs (Fig. 5B–F, H), we identified a vertebrate evolutionarily conserved 588 bp fragment - eR3-Lim, that was sufficient to mediate expression in commissural neurons along the dorsal–ventral axis (Fig. 5A, F, H). Co-staining with cell fate markers confirmed expression in commissural subtypes, with no expression in ipsilateral-projecting neurons such as dI3 (Fig. S7). Deletion of eR3-Lim from eR3-123 abolished reporter gene expression in 5 of 7 embryos, with only low-level nonspecific expression in 2 embryos (Fig. 5G). The mouse homologous eR3-Lim also drove reporter expression in commissural neurons in transgenic mice (Fig. S8B). We conclude that eR3-Lim is a commissural-specific enhancer required for Robo3 expression (Fig. 7I).

**Figure 5:**
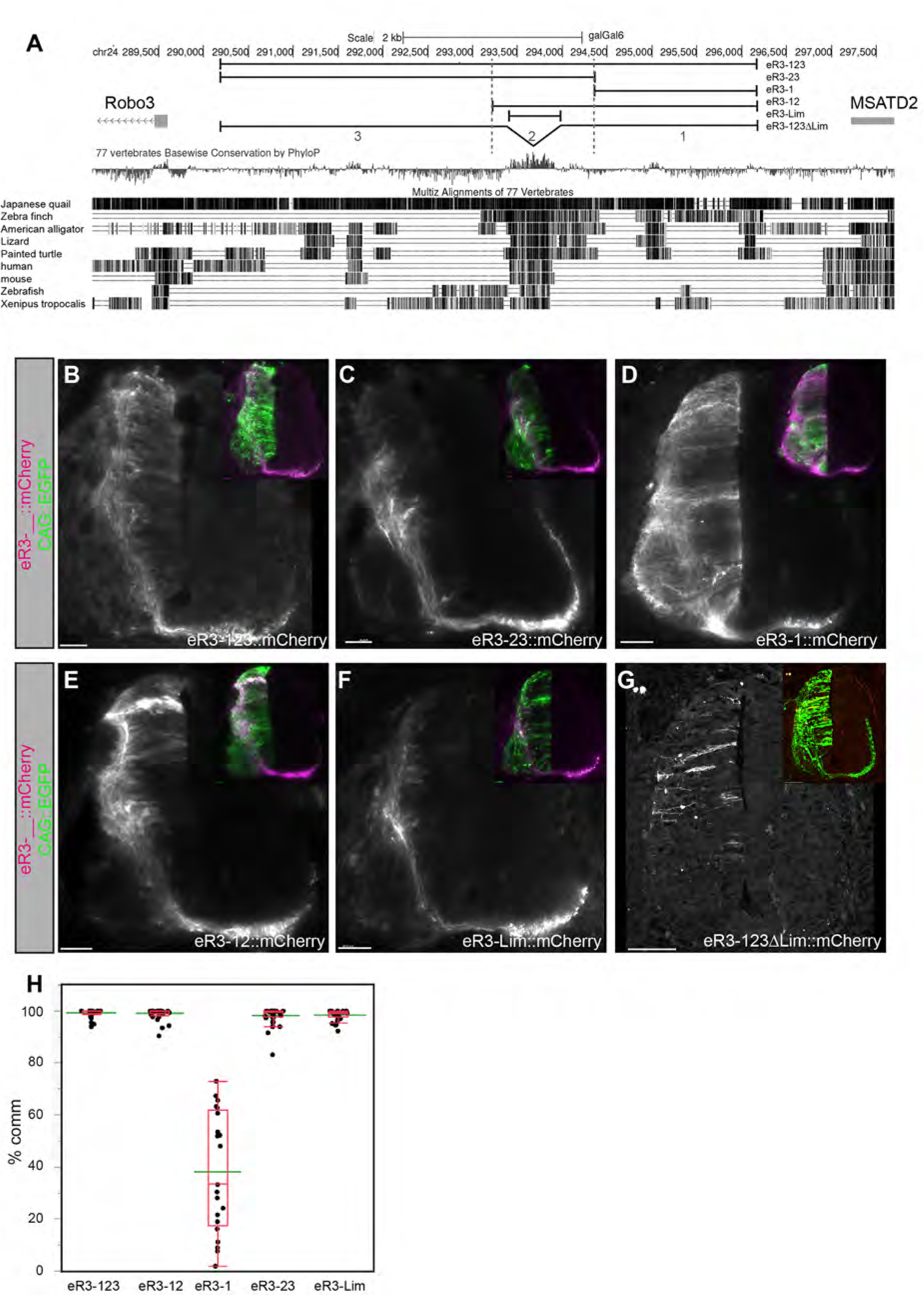
Identification of a Robo3 enhancer. A. Schematic of the *Robo3* enhancer region tested. The intragenic region between Robo3 and MSANTD2 of the chick genome. 5503bp were isolated (galGal6: chr24:290,319-295,821) and nested deletions were generated. The intragenic fragment is subdivided to three sub-fragments 1-3 (gray dashed lines). The nested deletion constructs are named accordingly: eR3-123: chr24:290,319-295,821; eR12: chr24:293,016-295,821, eR3-3: chr24:290,319-293,035; eR3-23: chr24:290,319-293,995; eR3-Lim: chr24:293,374-293,961. The fragments were cloned upstream to the TK promoter and mCherry reporter. B-G. Transverse section of the embryonic chick spinal cord showing activity of the enhancer constructs. The various eR3 enhancer, which drive expression the mCherry, were co-electroporated with CAG::EGFP. The expression of EGFP and mCherry is shown in the insets. H. Quantification of the mCherry commissural axons in the various enhancer constructs. (See supplementary statistics). Scale Bars = 50μm

Next, we identified evolutionarily conserved putative Lhx2/9 transcription factor binding sites (TFBSs) within eR3-Lim (Fig. S9). Specific expression in commissural axons was abolished when the Lhx2/9 TFBSs were mutated (eR3-123Limm::mCherry) (Fig. 6A, D), supporting the hypothesis that eR3-Lim is directly controlled by Lhx2/9. To test whether Lhx9 is sufficient to activate expression downstream of the Robo3 enhancer, we used both wild-type and a non-active mutant allele of Lhx9 (Lhx9m), which carries a point mutation in the homeodomain module (F315I) that abolishes DNA binding. Ubiquitous ectopic expression of Lhx9 with the eR3-123::mCherry reporter plasmid resulted in cell-autonomous reporter expression in both commissural and ipsilateral-projecting axons (Fig. 6C, D). The ectopic reporter expression in ipsilateral-projecting neurons reflects the lack of activation of the endogenous Robo3 (Fig. 4M). Expression of Lhx9m did not affect the commissural specificity of eR3-123 (Fig. 6B, D).

**Figure 6:**
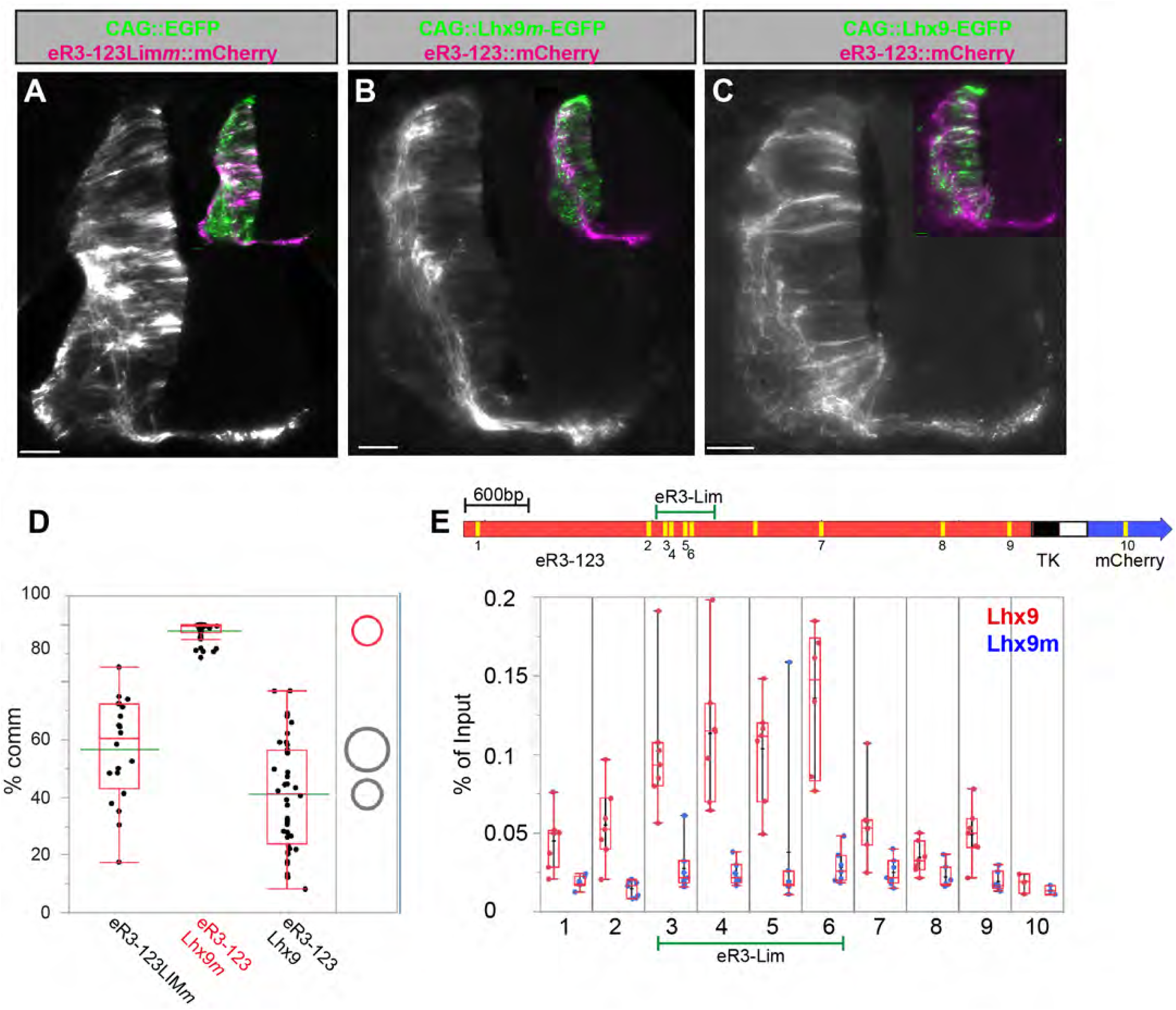
The core eR3-Lim element within the Robo3 enhancer is a target site for Lhx9. A-C: Transverse sections of E5.5 chick spinal cords co-electroporated with plasmid constructs of Robo3 enhancer (A-C), Lhx9m (B) and Lhx9 (C). A. The Lhx2/9 binding sites, within the core eR3-Lim domain were mutated. The mutated core element (eR3-Lim) is located within the Robo3 and MSANTD2 intragenic region – eR3-123Lim*m*. Ectopic expression of the mCherry reporter is apparent in many neurons as well ipsilateral and contralateral axonal projection. B. Co-expression of the mutated Lhx9*m* with eR3-123, results in exclusive commissural projection. C. Co-expression of the Lhx9 with eR3-123, results in ectopic expression of the mCherry reporter in many neurons as well ipsilateral and contralateral axonal projection. D. Quantification of the mCherry commissural axons in the various enhancer constructs. Comparing the ratio of commissural axons between the control: eR3-123+Lhx9*m*, and the experimental: eR3-123Lim*^m^* and eR3-123+Lhx9, using Dunnett’s method shows significant differences between the control in all experiments *P* < 0.05. (see Supplementary Statistics). E. Chip-PCR analysis of the *Robo3* and *MSANTD2* intragenic region. eR3-123::mCherry + Lhx9 or Lhx9*m* were transfected into COS cells. The binding of fragments 1-10 to either Lhx9 or Lhx9m was analyzed by PCR. Fragments 3-6 (See sequence in Fig. S9) are within the eR3-Lim core enhancer element. (see Supplementary Statistics). Scale Bars = 50μm

For loss-of-function experiments, we used a chimeric protein consisting of the DNA-binding homeobox domain of Lhx9 fused either to the repression domain of Engrailed (EnR-Lhx9) or to the repression domain of Barhl2 (BarEn-Lhx9), thereby converting Lhx9 from an activator into a repressor (Rodriguez-Esteban et al., 1998). In control experiments, using the mutant Lhx9m (En-Lhx9m and Bar-Lhx9m), the reporter gene was expressed in commissural axons. However, the proportion of commissural projections was significantly reduced following electroporation of En-Lhx9 or BarEn-Lhx9 (Fig. S10).

Direct interaction between transcription factors and TFBSs can be demonstrated by ChIP analysis. Analysis of ChIP-seq experiments with Lhx2 in the mouse retina (Zibes et al., 2019) and cortex (Ypsilanti et al., 2021) revealed Lhx2 binding sites within the eR3-Lim enhancer element (Fig. S8B). Next, we tested whether the minimal eR3-Lim, which includes the TFBSs, is a target of Lhx2/9. We performed ChIP-qPCR by co-transfecting eR3-123::mCherry with 6xmyc-tagged Lhx9 into COS cells. Lhx9m was used as a control. qPCR of four genomic regions within eR3-Lim (fragments 3–6) showed enrichment in the Lhx9-ChIP compared with the control Lhx9m-ChIP chromatin. By contrast, the five genomic regions flanking eR3-Lim (fragments 1, 2, 7–9) showed negligible enrichment (Fig. 6E). The ChIP-qPCR results demonstrate that eR3-Lim is an enhancer of Robo3, directly controlled by Lhx9.

Deletion of eR3-Lim abolished expression in all commissural neurons, not only in dI1 (Fig. 5G). Thus, we hypothesized that eR3-Lim may also serve as a target for other Lim-HD TFs in commissural neurons. The binding sites of Lhx2/9 and Lhx1/5 are highly similar (Fig. S9). To test whether the eR3-Lim enhancer is a target for Lhx1, we co-expressed Lhx1 with the eR3-Lim::mCherry enhancer plasmid. Co-expression with the core eR3-Lim enhancer induced ectopic expression (Fig. S11). Similar to Lhx2/9, Lhx1 did not induce expression of the endogenous Robo3 (Fig. S11). We conclude that Lhx1 activates expression through interaction with the eR3-Lim enhancer. The failure to induce endogenous Robo3 expression supports the assumption that Lhx1 activates the expression of a repressor (Fig. 7I).

**Figure 7:**
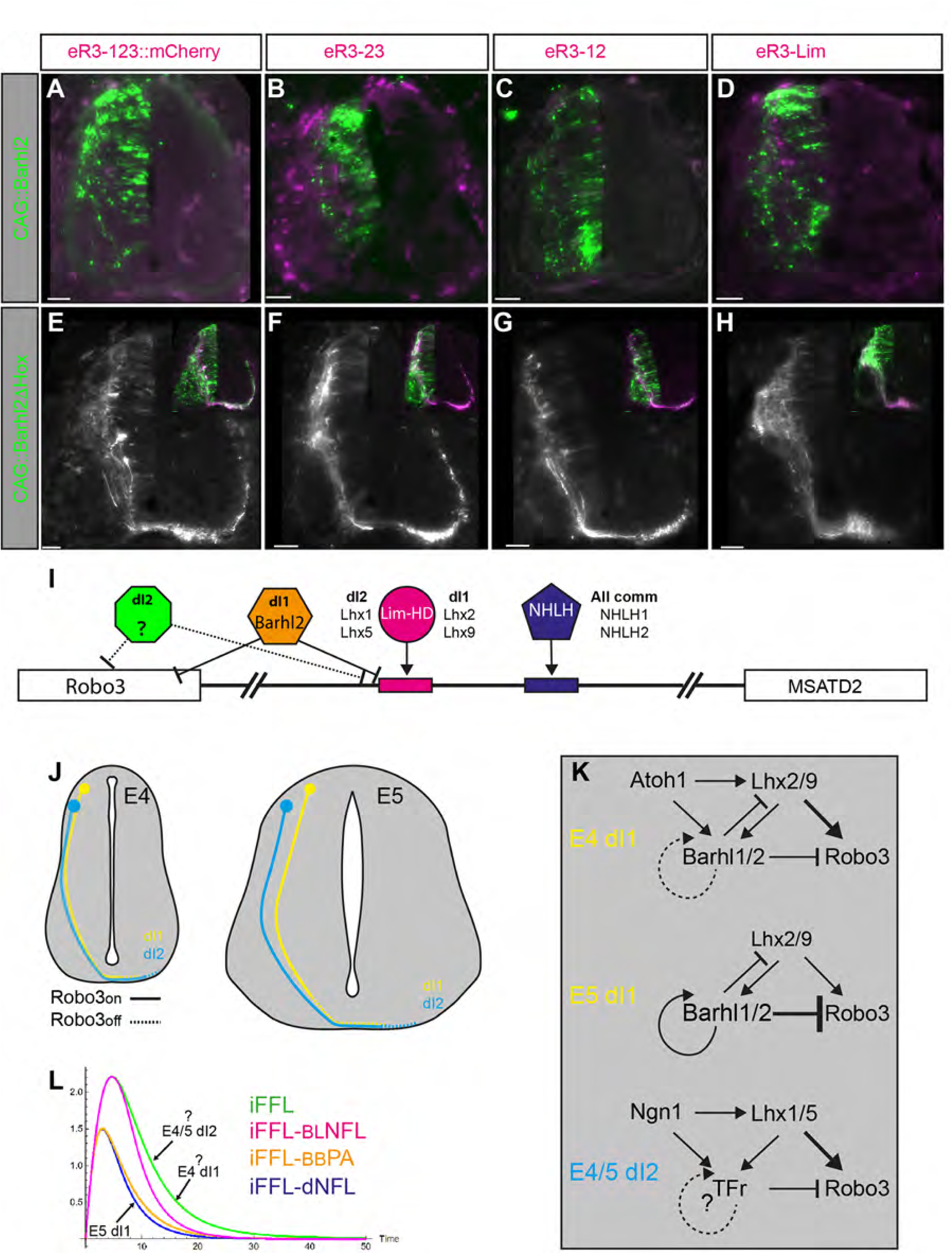
Barhl2 represses the enhancer activity of eR3-Lim. A-H. Transverse sections of E5.5 chick spinal cords co-electroporated with plasmid constructs of Robo3 enhancer with Barhl2-IRES-nEGFP (A-D) and Barhl21′home-IRES-nEGFP (F-H). Scale Bars = 50μm I. Schematic illustration of the iFFL’s TFs expressed in dI1 and in other commissural neurons (dI2), and the CIS control elements within the enhancers of Robo3. The broken lines indicate hypothetical GRNs. J. A scheme describing the presumed effect of the growth of the spinal cord, along the dorsal/ventral, on the laterally choice of E4 dI1 versus E5 dI1, and dI1 versus the all-comm cardinal subtype dI2. According to the model, at E5, the 20% increase of the longitudinal axis of the spinal cord, lengthen the trajectory path of the pre-crossing dI1 to the extent that expression of Robo3 is downregulated in pre-crossing axons. However, the path-extension does not affect dI2 since they are located more (full line – Robo3^+^, broken line – Robo^-^). K. Suggested iFFLs that explain the laterally choice of E4 dI1 versus E5 dI1, and dI1 versus dI2. At E4 dI1, the expression of Atho1 activator is balancing the repression by Barhl2, thus may eliminate the negative feedback inhibition of Barhl2 over Lhx2/9. The Barhl1/2 levels may not be sufficient for activating their own expression (broken arrowed curved line). Hence, the Barhl1/2 self-activation may also be compromise. Therefore, at E4 the expression of Robo3 might be controlled by a simple iFFL, as appose to the iFFL-dNFL in E5 dI1. The expression of Robo3 in dI2 may also be controlled by a simple iFFL. The presumed TFr may suppress Robo3 without self-activation and downregulation of Lhx1/5. L. The profile of Robo3 transient expression in E4 dI1, E5 dI1 and dI2 according to the suggested models in J and K.

### Barhl2 Represses the Activity of the Robo3 Enhancer

Ectopic expression of Lhx9 activated expression downstream of the exogenous eR3-Lim enhancer (Fig. 6C,D), while the endogenous Robo3 gene was not upregulated (Fig. 4M). It is plausible that Barhl2 mediates repression by binding to TFBSs outside the Robo3-MSANTD2 intragenic region (Fig. 7I).

To test whether the Robo3-MSANTD2 intragenic region and the eR3-Lim enhancer are subject to repression by Barhl2, we co-electroporated the enhancer reporter plasmids (Fig. 5A) with either Barhl2 or a mutated form of Barhl2 lacking the homeodomain (Barhl2ΔHox). Barhl2 repressed reporter gene expression in all reporter plasmids (Fig. 7A–D), whereas Barhl2ΔHox did not abolish commissural-specific expression (Fig. 7E–H). We conclude that Barhl2 can mediate repression by inhibiting the activity of the eR3-Lim enhancer. However, repression through other regulatory elements located outside the Robo3-MSANTD2 intragenic region also contributes to the inhibition of endogenous Robo3 expression (Fig. 7I).

## Discussion

Spatial and temporal precision in gene expression is critical for accurate embryonic development and stem cell differentiation in adults. Gene regulatory networks, such as coherent and incoherent feed-forward loops and feedback loops, play a significant role in controlling the onset, extent, and termination of gene expression. These networks regulate cellular responses and maintain homeostasis in both cells and organisms (Adler et al., 2014; Mangan and Alon, 2003; Mangan et al., 2006) and are likely fundamental to developmental processes. In the current study, we demonstrate that in the embryonic spinal cord, iFFL combined with feedback loops play a role in axon guidance. We reveal a neuronal-intrinsic iFFL that regulates the time frame of axonal attraction to the midline, determining whether an axon will cross to the contralateral side or remain on the ipsilateral side. This binary choice has implications for both motor and sensory physiology. The expression of *Robo3* is transcriptionally controlled by Lhx2/9 and Barhl2 via an iFFL. Lhx2/9 activate the expression of *Robo3* and the expression of the repressor *Barhl2*, which subsequently downregulates *Robo3*. The duration of attraction to the floor plate can be further modulated by strengthening the repression module of the iFFL through additional feedback loops that inhibit the activators (Lhx2/9) and stabilize the expression of the inhibitor (Barhl2). Our genomic analysis reveals that the iFFL converges on an enhancer element of *Robo3*. Together, our findings support the concept that a range of iFFLs control the timing and extent of midline crossing in commissural neurons of the spinal cord.

### Transition from Commissural to Ipsilateral Axonal Bias in Commissural Neurons

In adult mice, dI1 neurons consist of glutamatergic neurons that project predominantly ipsilaterally and are involved in regulating the frequency of locomotor output, suggesting that they play a role in rhythm generation alongside other spinal populations (Bertho et al., 2024). The transition from early commissural to late ipsilateral projections in the chick embryo is evolutionarily conserved and is also observed in mice (Wilson et al., 2008; Yuengert et al., 2015). Although most dI1 neurons shift to ipsilateral projections, in mice about 20% project contralaterally (Pop et al., 2022). It has been suggested that dI1_ipsi_ and dI1_comm_ neurons form subsets of the DSCT and VSCT, respectively (Yuengert et al., 2015).

The transition in axonal laterality bias, from contralateral to ipsilateral, is most apparent in the chick between E4.5 and E5 (Fig. 1A,B). During this period, the levels of Lhx2/9 and Barhl2 proteins are similarly upregulated (Fig. 1F, G, N, S1D). This pattern is also reflected in mRNA expression during the corresponding developmental stages in mice (Fig. S1E). The balance of TF levels shifts toward high Barhl2 and low Lhx2/9 at E6 in chicks and E12.5 in mice. This raises the question: what drives the switch in axonal laterality at E5 in chick and E11.5 in mouse embryos? We suggest two models that can explain the axonal laterality shift: 1. A change in spinal cord size. Between E4 and E5, the length of the dorsal/ventral axis of the spinal cord increases by 15–20% (Kicheva et al., 2014; Kuzmicz-Kowalska and Kicheva, 2021). It is plausible that the time frame of *Robo3* expression at E4 in chick embryos allows sufficient time for axons to reach and cross the floor plate. However, this time frame may be too limited for them to enter the floor plate at E5 (Fig. 7J). 2. A change in the type of iFFL. At earlier stages (E4 chick), the levels of Barhl2 may be insufficient to inhibit Lhx2/9 due to the expression of their activator Atoh1 and/or the low affinity of Barhl1/2 for the cis-regulatory elements that mediate their autoregulation. At E5, Atoh1 expression is downregulated, and Barhl1/2 rise above the threshold required for positive autoregulation (Fig. 7K,L). Consequently, *Robo3* is regulated by a simple iFFL at early stages, while at later stages it is regulated by an integrated mechanism of iFFL combined with feedback loops (iFFL + dNFL) (Fig. 7K,L). Our mathematical simulations suggest that the duration of *Robo3* expression is longer in a simple iFFL compared to the iFFL+dNFL combination (Fig. 4A–C). This may explain the earlier crossing of axons versus the later ipsilateral turning.

Similar hypotheses can explain the differences in axonal laterality choices between dI1 and other, more ventral commissural subtypes. While the laterality choice in dI1 is converted from commissural to ipsilateral, the commissural laterality of other commissural cardinal subtypes, such as dI2, V0, and V3, is predominantly retained (Alaynick et al., 2011; Lai et al., 2016). dI1 INs are located at the most dorsal position, while other commissural INs differentiate more ventrally. The shorter distance to the floor plate along the D/V axis may allow their axons to cross the floor plate within a shorter timeframe, before Robo3 expression is downregulated (Fig. 7J). Additionally, Robo3 regulation in the more ventral commissural INs may be mediated by a simple iFFL, in contrast to the iFFL+dNFL in dI1 (Fig. 7L). Future studies aimed at identifying the presumed Robo3 repressor in other commissural INs will help clarify the GRN that regulates Robo3 in these additional populations.

### Default Commissural Projection and Repression by Barhl2

We demonstrated that different iFFLs regulate a range of transient *Robo3* expression profiles that differ in their duration. Thus, dI1_ipsi_ neurons express *Robo3* for too short a period to enable floor plate crossing. However, we cannot exclude the possibility that the most restrictive GRN - iFFL+dNFL, prevents *Robo3* transcription entirely. At E5, the levels of Barhl2 may block *Robo3* transcription even in the presence of Lhx2/9. This hypothesis is supported by the commissural-only projection of Robo3^+^ neurons that stably express a reporter gene (Tulloch et al., 2019). It is plausible that the default axonal projection choice of dI1 neurons, governed by Lhx2/9 expression, is commissural, and that Barhl2 represses this default. Removal of Barhl2, as in null mice (Ding et al., 2012) or via expression of a dominant-negative isoform (Fig. 5), reveals the default commissural projection. A similar suppressed commissural-default may also account for the ipsilaterally projecting INs. Lim-HD as well as NHLH1/2 TFs are also expressed in the ipsilateral cardinal INs. Therefore, that binary choice of commissural versus ipsilateral axonal projection may account to the transient activation versus sustained repression of *Robo3* in commissural and ipsilateral projecting neurons, respectively.

## Materials and Methods

### Mathematical model of the gene regulatory network controlling Robo3 expression

#### Model of the i1FFL circuit with negative feedback on Lhx2/9 and positive autoregulation of Barhl1/2 (iFFL+dNFL)

To model the dynamical behavior of Robo3, we define a system of three coupled ordinary differential equations (ODEs) to describe the dynamical behavior of Lhx2/9 (ℓ), Barhl1/2 (b), and Robo3 (r). We assume that Lhx2/9 positively regulates Barhl1/2, while Barhl1/2 represses Lhx2/9. Additionally, Barhl1/2 exhibits self-activation, contributing to its sustained expression. Robo3, the terminal output of this circuit, is activated by Lhx2/9 and repressed by Barhl1/2. This configuration establishes a complex yet interpretable gene circuit, forming an incoherent feed-forward loop (iFFL) with feedback. The equations that describe this regulatory circuits thus reads:

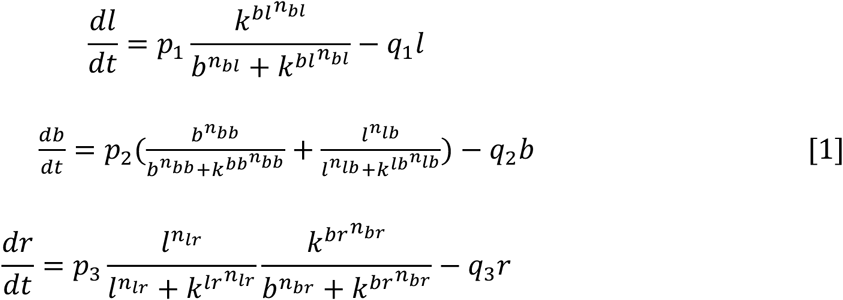

In these equations, l(t), b(t) and r(t) represent the concentrations of Lhx2/9, Barhl1/2 and Robo3, respectively. Each term uses Hill-type activation and repression functions, which are standard in modeling transcriptional regulation. Activating functions are written as 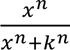 while repressive functions are modeled as 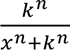, reflecting cooperative interactions and threshold behavior. Parameters *p_i_* are the maximum synthesis rates of Lhx2/9, Barhl1/2 and Robo3 and *q_i_* denote first-order degradation rates of each component. *k_i_*define Hill constants governing the strength of regulatory interactions and Hill constants governing the strength of regulatory interactions and *n_i_* are Hill coefficients, representing cooperativity in regulation indicating sensitivity of each regulatory effect.

The full circuit model successfully recapitulates key features of the experimentally observed midline-crossing regulatory circuit, including the mutual antagonism between Lhx2/9 and Barhl1/2, as well as the dual regulatory control of Robo3 expression. For a broad and biologically plausible range of parameter values (Table 1), the system dynamics governed by the combined incoherent feed-forward loop and negative feedback loop (<u>iFFL+dNFL</u>) converge to a steady state characterized by sustained activation of Barhl1/2 and decay of Lhx2/9. Under these conditions, Robo3 exhibits a transient pulse of activity in response to the temporal decline of Lhx2/9, followed by rapid repression driven by accumulated Barhl1/2 (Fig. 2 C, C’,C”). This behavior emerges robustly across parameter space, highlighting the circuit’s capacity to encode a temporally controlled pulse in Robo3 expression.

**Table 1:** List of parameters used in the mathematical modelling approach.

| Parameter | Biological meaning | Notes | Values | Units |
| --- | --- | --- | --- | --- |
| p<sub>1</sub> | Maximum synthesis rate of Lhx2/9 | Controls production strength of Lhx2/9 | 1 | a.u./time |
| p<sub>2</sub> | Maximum synthesis rate of Barhl1/2 | Controls production strength of Barhl1/2 | 1 | a.u./time |
| p<sub>3</sub> | Maximum synthesis rate of Robo3 | Controls production strength of Robo3 | 1 | a.u./time |
| $q_1$ | Degradation rate of Lhx2/9 | First-order decay constant | 0.4 | 1/time |
| $q_2$ | Degradation rate of Barhl1/2 | First-order decay constant | 0.01 | 1/time |
| $q_3$ | Degradation rate of Robo3 | First-order decay constant | 0.24 | 1/time |
| $k^{bl}$ | Hill constant for Barhl1/2 repression of Lhx2/9 | Repression threshold of Barhl1/2 on Lhx2/9 | 1.92 | a.u. |
| $k^{bb}$ | Hill constant for Barhl1/2 self-activation | Threshold for Barhl1/2 positive feedback | 0.1 | a.u. |
| $k^{lb}$ | Hill constant for Lhx2/9 activation of Barhl1/2 | Threshold for Lhx2/9–induced Barhl1/2 expression | 0.16 | a.u. |
| $k^{lr}$ | Hill constant for Lhx2/9 activation of Robo3 | Threshold for Lhx2/9–induced Robo3 expression | 0.18 | a.u. |
| $k^{br}$ | Hill constant for Barhl1/2 repression of Robo3 | Threshold for Barhl1/2–mediated repression | 5.32 | a.u. |
| $n_{bl}$ | Hill coefficient for Barhl1/2 repression of Lhx2/9 | Cooperativity of repression | 4 | dimensionless |
| $n_{bb}$ | Hill coefficient for Barhl1/2 self-activation | Cooperativity of positive feedback | 4 | dimensionless |
| $n_{lb}$ | Hill coefficient for Lhx2/9 activation of Barhl1/2 | Cooperativity of activation | 4 | dimensionless |
| $n_{lr}$ | Hill coefficient for regulation of Robo3 by Lhx2/9 | Cooperativity of activation | 4 | dimensionless |
| $n_{br}$ | Hill coefficient<br>for regulation<br>of Robo3 by<br>Barhl1/2 | Cooperativity<br>of repression | 3 | dimensionless |

**Table 2:**
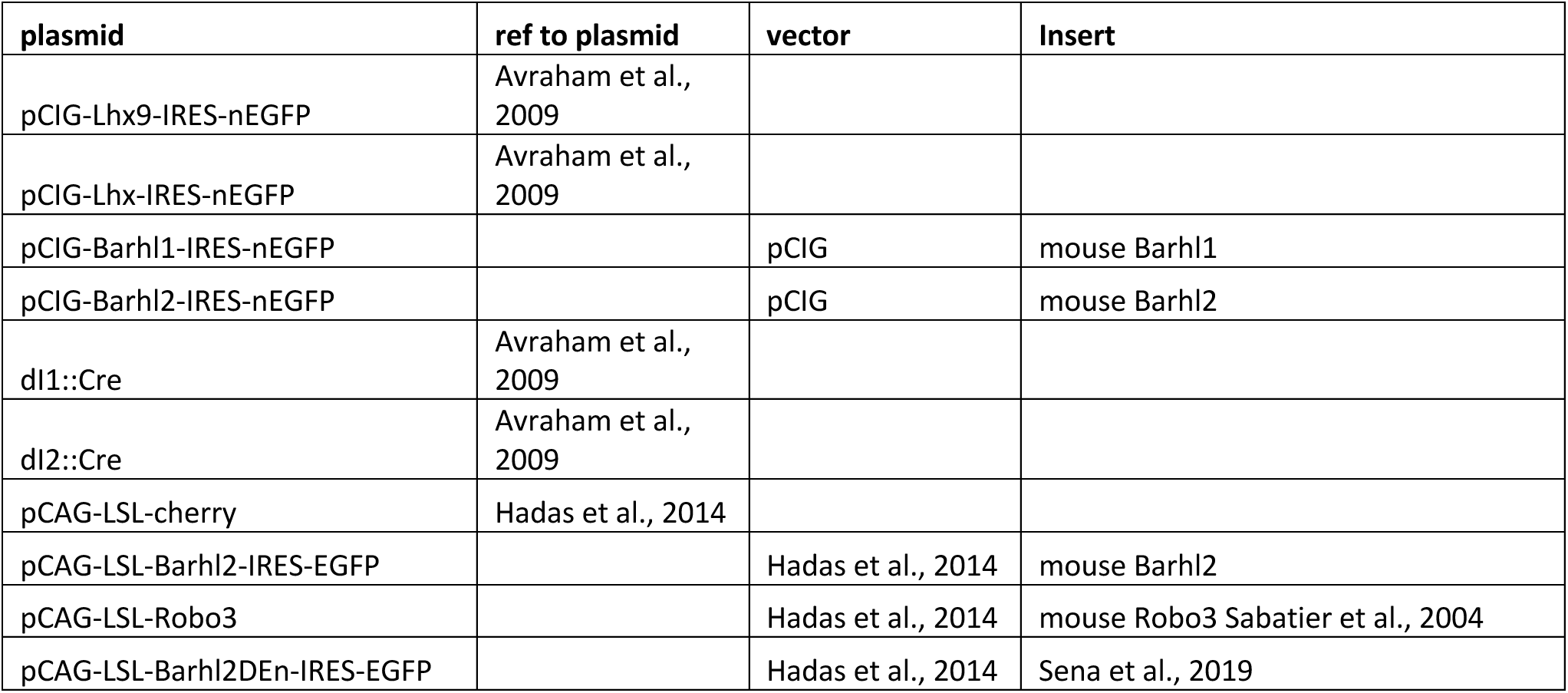
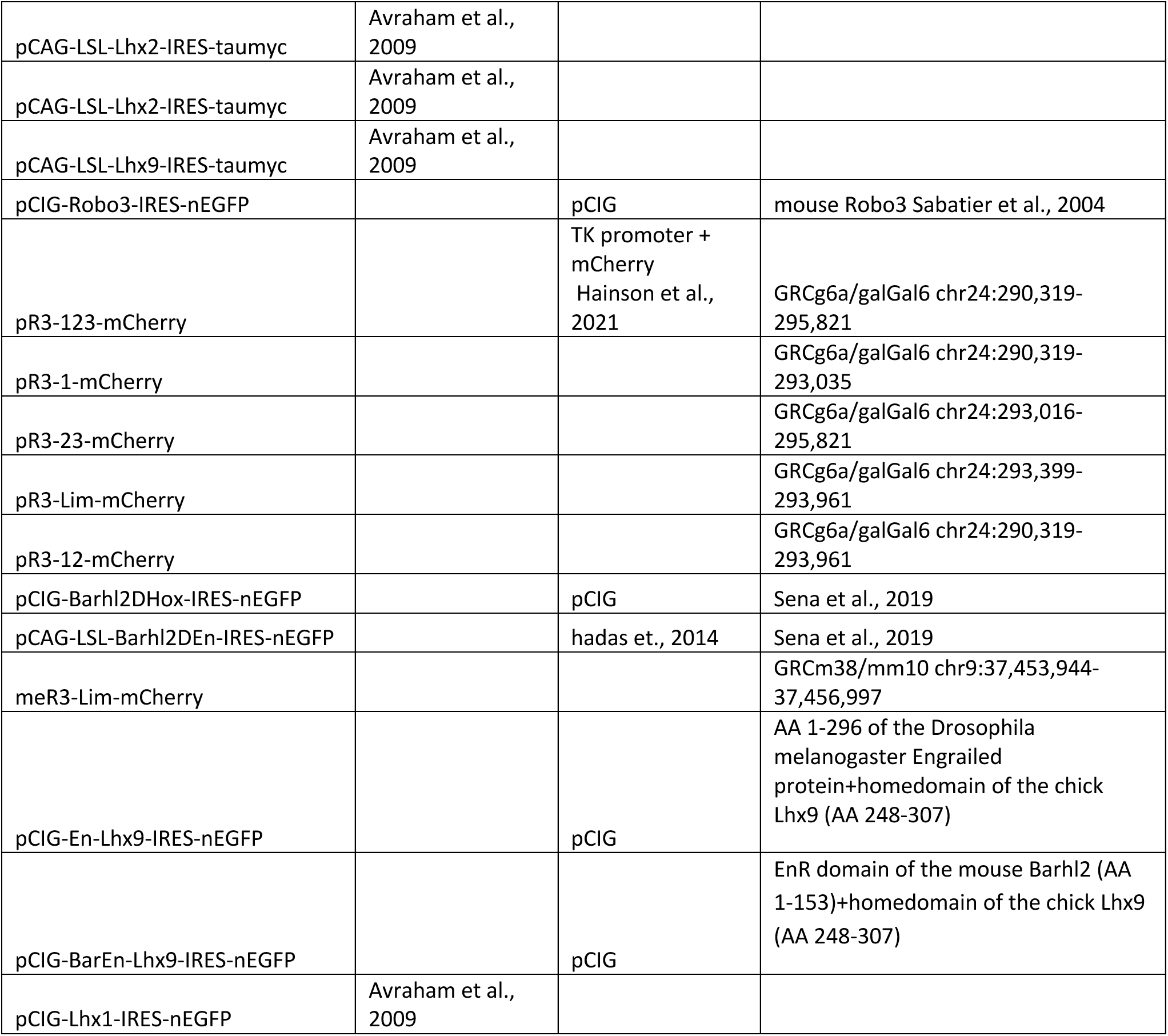
List of plasmids and the cloning strategy.

Next, we compare different variants of this regulatory circuit in the context of midline crossing to identify topological features that lead to optimal and robust behaviors. To investigate the role of circuit topology in shaping Robo3 dynamics, we systematically compared this canonical model to three simplified variants:

#### Minimal i1FFL without feedback or autoregulation (iFFL)

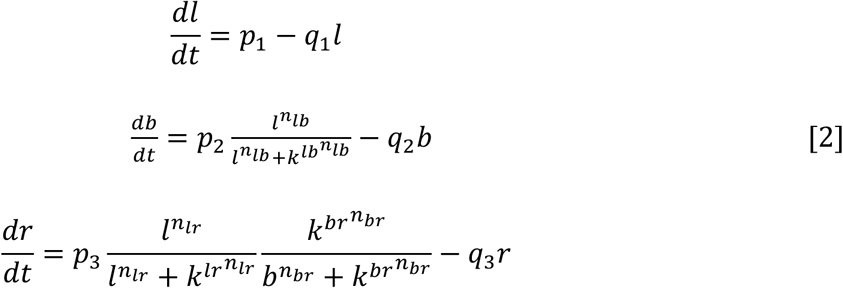

This minimal i1FFL model omits both Barhl1/2 self-activation and its inhibition of Lhx2/9. Here, Robo3 is activated directly by Lhx2/9 and repressed by Barhl1/2, which is itself induced by Lhx2/9. This topology is known to give rise to pulse-like outputs or perfect adaptation, depending on parameter values. Indeed, simulations show that Robo3 exhibits a transient expression profile (Fig. 2A), although with less precise timing and a broader temporal profile compared to the full circuit.

#### I1FFL with autoregulation but no feedback to Lhx2/9 (iFFL+BBPA)

We next investigated whether the simple incoherent feed-forward loop (<u>iFFL</u>) is sufficient to capture the pulse-like transient expression dynamics of Robo3 and questioned the necessity of Barhl1/2 autoregulation and its inhibitory feedback to Lhx2/9. To address this, we considered an alternative circuit configuration where Barhl1/2 maintains self-activation but no longer inhibits Lhx2/9. The dynamics of the system are governed by the following set of differential equations:

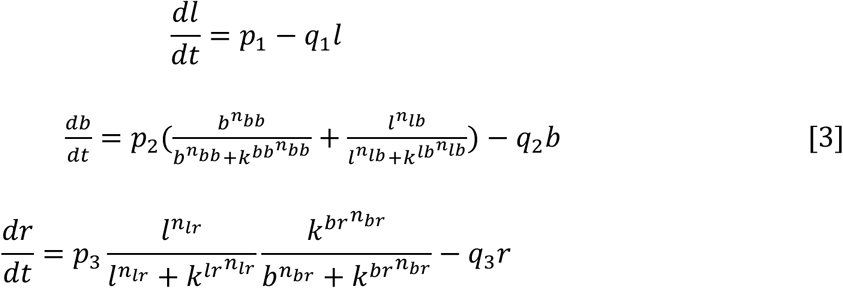

The first equation of this model (system 3) represents Lhx2/9 as a constitutively expressed factor undergoing linear degradation. In contrast to the full model (system 1), there is no inhibition from Barhl1/2 to Lhx2/9. The phase portrait derived from the first and third equations of this model (system 3) demonstrates that the system trajectories converge to a steady state characterized by sustained activation of Lhx2/9 and repression of Robo3, indicating that in the absence of feedback inhibition, Lhx2/9 remains persistently active (Fig. 4 A, B, C). Despite this, Robo3 initially responds with a transient peak before returning to its baseline level. Consistently, the time series analysis reveals that Robo3 exhibits a pulse-like transient expression with a broader temporal width compared to the <u>iFFL+dNFL</u> circuit (system 1), reflecting the absence of rapid repression, in contrast to the maintained high levels of Lhx2/9 (Fig 4 A, B, C). Thus, the negative feedback from Barhl1/2 to Lhx2/9 serves a dual mathematical and biological function as it enables transient Robo3 expression by allowing Barhl1/2 to eliminate the activating input (Lhx2/9) and it shortens the width of the pulse by accelerating the decline of Lhx2/9 ensuring that Robo3 activation is precisely timed and sharply terminated, consistent with experimental observations of midline crossing (Fig. S6).

#### I1FFL with feedback but no autoregulation (iFFL+BLNFL)

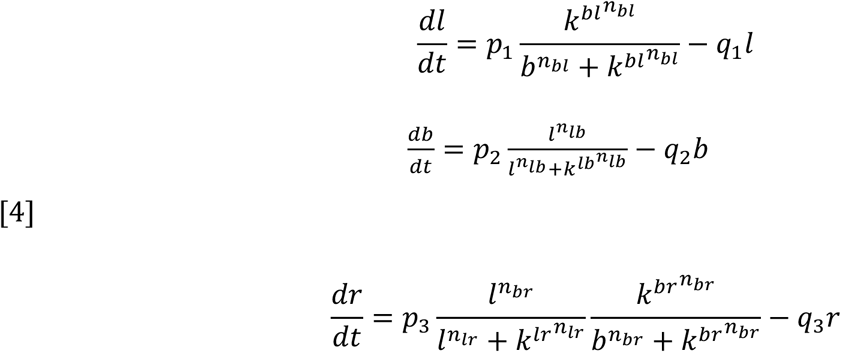

This model incorporates negative feedback from Barhl1/2 to Lhx2/9 but removes Barhl1/2’s self-activation. This configuration simplifies the downstream node while retaining mutual antagonism between the transcription factors. It preserves regulatory feedback without allowing Barhl1/2 to sustain its expression independently, possibly making the circuit more responsive and sensitive to changes in upstream Lhx2/9 activity. The time series plot illustrates the transient response of Robo3 for this model, but with a larger width of the pulse than the iFFL+dNFL (system 1). Ultimately this suggests incorporating positive autoregulation of Barhl1/2 enhances control over the amplitude and timing of the Robo3 response.

### Animal Models

#### Chicken

Fertilized white leghorn chicken eggs (Gil-Guy Farm, Israel) were incubated under standard conditions at 38°C. All experiments involving animals followed the designated policies of the Experiments in Animals Ethics Committee and were performed with its approval.

#### Mice

All experimental procedures had approval through Brown University’s Institutional Animal Care and Use Committee and followed the guidelines provided by the National Institutes of Health. Mice were maintained on a CD-1 background. For timed pregnancies, the day of vaginal plug was defined as embryonic day 0.5, and embryos of either sex were used for experiments.

### Mouse whole embryo culture

Spinal cord electroporation and *ex utero* culture of E9.5 embryos was essentially carried out as previously described (Chen et al., 2008). In short, equimolar amounts of an actin::GFP expression vector and the meLim::mCherry reporter construct (combined concentration: 0.1 mg/ml) were injected into the spinal cord central canal of E9.5 mouse embryos, followed by unilateral electroporation. Embryos were grown for 2 days at 37C in rat serum (Envigo/Inotiv) containing 10 mM glucose and 1 x Penicillin/Streptomycin/Glutamine (Invitrogen) under 60% O_2_/5% CO_2_ (day 1), followed by 95% O_2_/5% CO_2_ (day 2). Embryos were fixed in 4% PFA for 1 h, washed 3 x 10 min in PBS, cryoprotected O/N at 4C in 30% sucrose in PBS, embedded in OCT, and transverse cryosections were cut at 20 microns. Sections were directly mounted under Fluoromount G. Images were acquired on a Nikon Ti-E microscope.

### Chicken electroporation Procedure

In ovo electroporation was performed as previously described (Avraham et al., 2009). Briefly, fertilized chicken eggs were incubated until the desired developmental stage (typically HH stage 17-18 for E2.75-E3). A DNA solution (5 mg/mL) was injected into the lumen of the neural tube using a fine glass micropipette. Electroporation was carried out using three 50 ms pulses of 25-30 V applied across the embryo using 0.5 mm tungsten wire electrodes and a BTX electroporator (ECM 830).

Following electroporation, eggs were sealed and returned to the incubator. Embryos were harvested 2-4 days post-electroporation (at E5.5-E7.5) depending on the experimental requirements. Successful electroporation was confirmed by examining reporter gene expression (GFP or mCherry) in the targeted region.

### Molecular Biology and Gene Expression Constructs

Expression constructs were generated using standard molecular cloning techniques. Transcription factor coding sequences for Lhx2, Lhx9, Barhl1, and Barhl2 were cloned into appropriate expression vectors under the control of ubiquitous promoters to enable ectopic expression following electroporation (Hadas et al., 2014). Functional domain mutants of Barhl2 were obtained from Béatrice Durand (Sena et al., 2019). The mouse and chick cR3 enhancer elements were cloned by PCR from genomic DNA utilizing sequence-specific primers. The mutated cR3-Lim sequence was ordered as gBlocks DNA fragments (IDT).

### Analysis of Barhl2 expression in Lhx2 Lhx9 double null mice

The *Lhx2;Lhx9* mouse embryo tissue was generated in Professor Jane Dodds lab, Columbia University under approval by the IACUC and gifted for reuse. 3 E11.5 *Lhx2^+/-^;Lhx9^+/-^* and 9 control embryos were analyzed. Controls consisted of embryos which were either wildtype, single or double heterozygote for the *Lhx2* and *Lhx9* genes. *Math1LacZ* transgene was present in some embryos. Transverse spinal cord sections were labeled by immunohistochemistry with Barhl2 and DAPI and images acquired at subthreshold intensity. The raw images were analyzed in Cell Profiler (www.cellprofiler.org) where ‘objects’ (Barhl2^+^ cells) were delineated and the pixel count in each Barhl2^+^ cell quantified. This resulted in a pixel distribution of Barhl2^+^ cells in the spinal cord of control and double mutant embryos. As three different litters cohorts were analyzed, the data was normalized to control values within the same litter in order that data acquired form separate litter cohorts could be accurately compared. The data was statistically analyzed using Prism 9. Raw data pixel counts are available on request.

### Tissue Preparation

Embryos were fixed overnight at 4°C in 4% paraformaldehyde in 0.1 M phosphate buffer, followed by cryoprotection in 30% sucrose/PBS for 24 hours. Tissues were embedded in OCT compound (Scigen, Grandad, USA) and sectioned at 20 μm thickness using a cryostat. Sections were collected on Superfrost Plus slides and stored at −20°C until processing.

### Immunohistochemistry

Immunohistochemical analysis was performed using standard protocols. Primary antibodies included: rabbit anti-GFP (1:1000, Molecular Probes), mouse anti-GFP (1:200), goat anti-GFP (1:300, Abcam), rabbit anti-RFP (1:1000, Acris), mouse anti-Myc (1:10, 9E10), rabbit anti-Myc (1:100, Abcam), rabbit anti-HA (1:100, Abcam), rabbit anti-Barhl1 (1:200, Abcam), rabbit anti-Barhl2 (1:200, Novus Biologicals), goat anti-Lhx2/9 (1:100), and mouse anti-Lhx1/5 (1:100, 4F2, Hybridoma Bank, University of Iowa, USA). Secondary antibodies were Alexa Fluor 488/647-conjugated or Rhodamine Red X-conjugated donkey anti-mouse, anti-rabbit, and anti-goat antibodies (Jackson ImmunoResearch).

Sections were incubated with primary antibodies overnight at 4°C, followed by appropriate secondary antibodies for 2 hours at room temperature. Images were captured using either a fluorescence microscope (Eclipse Ni; Nikon) with a digital camera (Zyla sCMOS; Andor) or a confocal microscope (FV1000; Olympus).

### In Situ Hybridization

In situ hybridization was performed to detect Robo3 mRNA expression following gain-of-function experiments, particularly after Barhl2 overexpression. Antisense RNA probes were generated using standard protocols. Hybridization was carried out overnight at 65°C, followed by stringent washes and detection using alkaline phosphatase-conjugated anti-DIG antibodies.

### Quantification of commissural projection

Images of spinal cord cross sections were analyzed using ImageJ tools. Specifically, the number of pixels at the white matter on the ipsilateral and the contralateral spinal cord (without the floor plate) was scored. % comm = #commissural pixels / (#commissural pixels + ipsilateral pixles).

### Cell Culture and Transfection

COS-7 cells were cultured in Dulbecco’s modified Eagle’s medium (Sigma-Aldrich) supplemented with 10% fetal bovine serum, penicillin (100 U/ml), streptomycin (100 μg/ml), and 2 mM glutamine (Biological Industries). Cells were plated in 100 mm dishes at approximately 80% confluence and transiently transfected for 2 days with a mixture of 10 μg plasmid DNA (enhancer and cDNA plasmids in 1:1 ratio) using 25 μg polyethylenimine (linear, MW 25,000; Polysciences Inc.). For functional analysis of Barhl2 DNA-binding requirements, cells were also transfected with Barhl2ΔHD mutant constructs to serve as negative controls in ChIP assays, confirming the specificity of Barhl2-enhancer interactions.

### Chromatin Immunoprecipitation (ChIP) Assay

ChIP assays were performed to demonstrate direct binding of transcription factors to the Robo3 enhancer elements. Cells were harvested by scraping from 2 dishes, centrifuged at 600g for 5 minutes, and resuspended in 1 ml PBS. Cross-linking was performed by adding 100 μl of 11% formaldehyde solution for 15 minutes (Lhx9) or 5 minutes (Barhl2), then terminated with 60 μl of 2.5 M glycine solution.

Cell pellets were lysed in 300 μl RIPA buffer (10 mM Tris pH 7.4, 0.5 M NaCl, 1 mM EDTA, 1% Triton X-100, 0.1% sodium deoxycholate, 0.5% SDS, plus protease inhibitors) for 30 minutes on ice. Chromatin was sheared using a Bioruptor PLUS sonicator (2 rounds of 20 cycles, 30 sec ON/30 sec OFF at high power).

Immunoprecipitation was performed overnight at 4°C using 5 μg goat anti-Myc antibody (ChIP grade, ab9132, Abcam) or 3 μg rabbit anti-HA antibody (ChIP grade, ab9110, Abcam) coupled to 70 μl protein G magnetic beads (Dynabeads, Invitrogen). After extensive washing, DNA-protein complexes were eluted and cross-links reversed by incubation at 65°C for 4 hours in Direct Elution Buffer containing RNase A, followed by proteinase K treatment at 55°C for 2 hours.

### Real-Time qPCR Analysis

Immunoprecipitated DNA fragments were quantified using real-time PCR with primers encompassing different regions of the Robo3 enhancer (cAJ5-3) and control regions. PCR was performed using PerfeCTa SYBR Green FastMix (Quantabio) according to manufacturer’s instructions. ChIP enrichment was calculated using the delta Ct method and expressed as percentage of input.

### Image Processing and Documentation

All images were processed using ImageJ software for quantitative analysis and Adobe Photoshop for figure preparation. Confocal images were processed using manufacturer-provided software, with identical settings applied to experimental and control samples for comparative analysis.

### ChIP Seq Analysis

ChIP Seq data utilized in this study were previously published (Ypsilanti et al., 2021; Zibetti et al., 2019) and are accessible through GEO Series: GSE183130 (https://www.ncbi.nlm.nih.gov/geo/query/acc.cgi?acc=GSE183130), and GSE99818 https://www.ncbi.nlm.nih.gov/geo/query/acc.cgi?acc=GSE99818

Adapters were trimmed using the Cutadapt tool (version 2.7) (Martin, 2011). Following adapter removal, reads that were shorter than 30 nucleotides were discarded (Cutadapt option –m 30). The reads were aligned uniquely to the mouse genome (mm10) using Bowtie (v1.2.3)

(Langmead et al., 2009). Peaks were called using MACS2 using the options --SPMR to normalize the output signal (v2.0.10.20131216) (Zhang et al., 2008). MACS2 bedGraph files were converted to bigWig format for visualization using IGV (Thorvaldsdottir et al., 2013).

### Single Cell Data

Single-cell RNA-seq data utilized in this study were previously published (Delile et al., 2019; Rekler et al., 2024) and are accessible through GEO Series GSE261603 (quail) in ArrayExpress under accession number E-MTAB-7320 and (mouse).

The processing and analysis of the quail data is described in (Rekler et al., 2025; Rekler et al., 2024). For the analysis of the mouse single cell RNA-seq data,we excluded cells with total UMI counts or total expressed genes with more than 4 MAD defined by an adaptive thresholding method. Cells were also filtered using a 4 MAD cutoff for mitochondrial content. In addition, genes detected in fewer than ten cells were excluded from the analysis. The Seurat R package (Satija et al., 2015) (v4.9.9) was used for downstream analysis and visualization. We focused on day E11.5, that includes 372 cells, across 12,278 genes. The expression data was normalized and log-transformed using Seurat’s ‘NormalizeData’ function. The top 500 highly variable genes were identified using Seurat’s ‘FindVariableFeatures’ function with the ‘vst’ method. A potential source of unspecific variation in the data was removed by regressing out the UMI count. The data was scaled and centered as implemented in the function ‘ScaleData’ of the Seurat package. For PCA, we selected 25 principal components (PCs) for downstream analyses. Cell clusters were generated using Seurat’s unsupervised graph-based clustering functions ‘FindNeighbors’ and ‘FindClusters’ (resolution=0.1).

Seurat’s functions ‘FeaturePlot’ and ‘DimPlot’ were used for visualization. Plots were further formatted using custom R scripts with ggplot2. Heatmaps were produced with The R package ComplexHeatmap (Version 2.14) (Gu, 2022).

Trajectory inference and pseudotime estimation were performed using Slingshot (v2.7.0) (Street et al., 2018).

## Supporting information

Supplementary figures, statistics and Mathema3cal Analysis

## ACKNOWLEDGMENTS

The authors wish to thank Noa Tal, Roni Avigdory, Netanel Wote and Y. Zhou for expert technical assistance; Arthur Radley and James Briscoe for providing the E11.5 mouse scRNA data; Béatrice Durand for the Barhl2 mutant plasmids; Alain Chédotal and Yan Zhu for critical reading of the manuscript. The *Lhx2;Lhx9* mouse embryo tissue was generated in Professor Jane Dodds lab, Columbia university under approval by the IACUC and gifted for reuse.

## Funding

This work was supported by grants to: A.K. from the Israel Science Foundation (grant no. 1787/21) and the Avraham and Ida Baruch endowment fund; to M.A. from the Israel Science Foundation (ISF, Grant No. 3372/24), to A.J. from the National Institutes of Health (R01NS095908) and Carney Institute for Brain Science Zimmerman Innovation Award; to S.I.W: Lennart Glans stiftelse, Cancerfonden (24 3790 Pj), Cancerforskningfonden (AMP 24-1154), Kempestiftelsena (JCSMK25-0048, and JCK-1927, U17).

G.F. is the Incumbent of the David and Stacey Cynamon Research fellow Chair in Genetics and Personalized Medicine at the Weizmann Institute of Science.

## AUTHOR CONTRIBUTIONS

R.S.M, AK, and MA conceived the study. R.S.M, M.M, G.E., M.B., S.K., S.W.I. and A.J. performed experiments, R.S.M., M.M., G.E., G.F., A.J., M.A. and A.K. analyzed the data, D.N. and C.K. provided reagents and data, R.S.M, M.M., S.I.W., A.J., M.A. and A.K. interpreted data, R.S.M., M.M., M.A. and A.K. wrote the manuscript, with feedback from all co-authors.

