## Supplementary figures, statistics and Mathema3cal Analysis for "Temporal control of axonal floor plate crossing through a combination of incoherent feedforward and feedback loops of gene regulatory network regulating Robo3 expression"

Fig. S1. Dynamics and levels of TFs and Robo3 expression in mouse and quail dI1.

A. UMAP of E4 quail dorsal interneurons (Rekler et al., 2025; Rekler et al., 2024), colored by cluster identity. Cluster numbers are indicated both in parentheses (and displayed on top of each cluster). The fitted principal curves of dI1 and dI2 are shown in black.

B,C. Cells in the UMAP are colored by dI1 pseudotime (B) or dI2 pseudotime (C), as inferred by Slingshot (See Methods).

D,E. Expression profiles of selected genes along the final segment of the dI1 trajectory in Quail (D- clusters 9 and 4) (Rekler et al., 2024) and mouse (E) (Delile et al., 2019). Scaled expression values (computed using Seurat's ScaleData function) are plotted against the normalized pseudotime (scaled to the 0–1 range) of the dI1 trajectory. Points represent individual cells, and the black curves indicate LOESS-smoothed scaled expression values.

Figure S2. Barhl2 is downregulated in *Lhx2$^{-/-}$;Lhx9$^{-/-}$* double mutant mouse embryos.

**A-C**. Photomicrographs of transverse sections of control (**A,B**) and *Lhx2$^{-/-}$;Lhx9$^{-/-}$* double mutant (**C**) mouse embryos labeled immunofluorescently with Barhl2. (**D-F**) Quantification of the Barhl2 pixels count in 9 control and 3 *Lhx2$^{-/-}$;Lhx9$^{-/-}$* double mutant mouse embryos. (**D**) A graph depicting the number of Barhl2+ cells in control and *Lhx2$^{-/-}$;Lhx9$^{-/-}$* double mutant embryos. An unpaired t-test was used. The normalized Barhl2$^{+}$ pixel count was quantified for each cell and Barhl2+ cell in control and *Lhx2$^{-/-}$;Lhx9$^{-/-}$* double mutant embryos compared for all cells using One sample t and Wilcoxon test (**E**) or by comparing as a mean Barhl2+ pixel count per embryos using an unpaired t test (**F**). on the graphs ns, ** and **** are not significant (**D**), P<0.0045 (**F**) and P<0.0001 (**E**) respectively. DKO = *Lhx2$^{-/-}$;Lhx9$^{-/-}$* double mutant embryo.

Fig. S3. Ectopic Lhx2 and Lhx9 rescue the ectopic Barhl2-induced ipsilateral bias of axonal projection.

Transverse sections of chick E5.5 spinal cords are shown. The following plasmids were electroporated: dI1::Cre (A-D), CAG::LSL-Barhl2-IRES-EGFP (A-D), CAG::LSL-Lhx2-IRES-taumyc (B), CAG::LSL-Lhx9-IRES-taumyc (C,D). Three examples from each electroporation are shown. The inset on C'' is shown in D.

A. Ectopic expression of Barhl2 induces ipsilateral turning (Magenta arrows).

B. Ectopic expression of Barhl2+Lhx2 restores commissural trajectory (Yellow arrows).

C. Ectopic expression of Barhl2+Lhx9 restores commissural trajectory (Yellow arrows).

D. Axon co-expressing Barhl2+Lhx9 express Robo3 (Yellow arrow).

Fig. S4 Ectopic expression of Lhx2/9 in dI1 impose a transient ipsilateral axonal deflection at E5.

A-D. Transverse sections of the E5.5 chick spinal cords are shown. Ectopic expression of Lhx2 or/and Lhx9 in dI1. The following combinations of plasmids was used: dI1::Cre (Avraham et al., 2009) (A-D), CAG::LSL-EGFP (A) ,CAG::LSL-Lhx2-IRES-taumyc (B,D), CAG::LSL-Lhx9-IRES-EGFP (C,D). Scale Bar = 50$\mu$m

E. Quantification of extent of ipsilateral turning in dI1 following Lhx2/9 expression in dI1 at E5. The average % of commissural axons from the total axons is: 60.74+/-11% of the EGFP-expressing dI1. In dI1 neurons expressing Lhx2 (46.98+/-13%), Lhx9 (45.82+/-11%) or Lhx2+9 (40.96+/-15%). Comparing the % of ipsilateral axonal projection between control-EGFP to the experimental Lhx2/9 across groups using Dunnett's method (which takes into account multiple comparisons) shows significant differences between the EGFP and the Lhx2/9 expressing dI1 neurons in all experiments. The circle charts show the significance. No overlapping between the experimental electroporated side (gray circles) and control-nonelectroporates (red circle) is indicative of $P < 0.05$ (see Supplementary Statistics).

Fig. S5. Ectopic expression of Lhx1 does not upregulate the expression of Lhx1.

Transverse section of the E5.5 chick spinal cord following ectopic expression of CAG::Lhx1-IRES-nEGFP. The pattern of Lhx1-expressing neurons (left side of the spinal cord) does not change following ectopic expression of Barhl2.

Fig. S6. Impact of network topology and feedback strength on Robo3 pulse dynamics and phase space trajectories.

A. Time-course profiles of simulations of Robo3 expression across four different network topologies, as illustrated on the right. Each topology differs in the presence or absence of feedback interactions involving Barhl1/2 and Lhx2/9 or the autoregulation of Barhl1/2, and curves are color-coded accordingly. Despite all topologies producing a transient Robo3 response, the rate of decline varies with network configuration, with negative feedback accelerating signal decay.

B. Phase portraits in the Lhx2/9–Robo3 space under two different conditions. Blue lines denote Lhx2/9 nullclines (lines in which Lhx2/9 levels do not change over time) and magenta trajectories show Robo3 nullclines for decreasing levels of Barhl1/2, as shown in the figure. Points where the blue and magenta lines intersect mark the levels of Robo3 as function of time. Red dot marks the peak of Robo3 response, and the black dots mark the final steady state level of Robo3.

Left panel: The system with only repression of Robo3 by Barhl1/2 without inhibitory feedback to Lhx2/9 (simple i1FFL). Robo3 level converge slowly toward the steady state (black dot), with decay speed modulated by $\frac{1}{b^{n_{br}}}$.

Right panel: The system incorporating negative feedback from Barhl1/2 to Lhx2/9, in addition to Robo3 repression (i1FFL+NFL). The combined influence of $1/b^{n_{br}}$ and $1/b^{n_{bl}}$, results in a faster approach to the steady state and earlier shutdown of Robo3 expression.

Fig. S7. The Robo3 enhancers drives expression in various cardinal subtypes of spinal interneurons.

The chick and mouse eLim Robo3 enhancers, driving expression of reporter genes, were electroporated into the chick spinal cord. Expression in cardinal subtypes of spinal INs was tested by using cell fate markers.

A-C. The chick enhancer drives expression in the commissural V0 ($Evx1^+$ neurons, A), dI2 (the dorsal $Lhx1^+$ and the more ventral $Lhx1^+$ neurons, B) and dI4 INs, but not in the ipsilateral dI3 ($Isl1^+$ neurons, C). The mouse enhancer drives expression in dI1 (Barhl1+ neurons, D), dI2 and dI4 (Lhx1+ neurons, E), but not in the ipsilateral dI3 ($Isl1^+$ neurons, F).

Fig. S8. Genomic organization of Robo3's enhancers in the mouse genome.

A. Genomic organization of the mouse Robo3-Msantd2 intragenic region. The eR3-bHLH (Masuda et al., 2024) and the eR3-Lim (this study) enhancers are situated 5' to Robo3. The eR3-Lim includes four ENCODE cCRE; are in open chromatin configuration in the embryonic brain, as assessed by ATAC seq; associated with the Robo3 enhancer; and conserved between vertebrates.

B. Analysis of Chip-seq data of Lhx2 in the retina (Zibetti et al., 2019) and the cortex (Ypsilanti et al., 2021), revealed two statistically significant peaks within eR3-Lim (red arrows). The 5' peak (red rectangular chr9:37,454,076-37,454,504) is homologous to the chick eR3-Lim core element. IGV visualization of normalized ChIP Seq signal (bigWig files generated by MACS2) for both ChIP and control samples. The peaks highlighted in red arrows were consistently identified by MACS2 across all samples.

C. A 3054bp genomic mouse DNA, which includes the four ENCODE cCREs within eR3-Lim (A) was clone upstream to the TK promoter and mCherry (meR3-Lim::mCherry) and co-electroporated with CAG::GFP into E10 mouse spinal cord. The embryos were cultured in vitro and analyzed at E11.5. meR3-Lim drives expression in commissural axons.

D. The percentage of commissural axons expressing EGFP under the control of the ubiquitously expressed CAG enhancer, and mCherry under the control of the eR3-Lim enhancer, was analyzed in the same cross-section. The average percentage of commissural projections in six embryos is presented. A relative enrichment of commissural projections expressing eR3-Lim::mCherry compared to those expressing CAG::EGFP is observed in all six embryos. For more details, see supplementary statistics.

Fig. S9. Sequence and genomic conservation of the eR3-Lim enhancer.

A. Lhx2/9 and Lhx1/5 transcription factor binding sites (TFBSs) according to JASPAR data of transcription factors binding sites (https://jaspar.elixir.no/).

B. Sequence and alignment of the eR3-Lim enhancer of human, mouse, opossum, chicken, zebra finch, America alligator and xenopus tropicalus. The Lim-HD TFBSs are marked in green and the mutations that we generated in red. The colored arrows mark the PCR fragments, within the core eR3-Lim enhancer, that we used in the Chip-PCR assay (Fig. 6E).

Fig. S10. Dominant negative Lhx9 induces ipsilateral axonal projection of commissural axon.

The Engrails domains of the drosophila Engrailed (A,B) or Barhl2 (D,E) were fused to the homedomain of Lhx9 (A,D) or Lhx9*m* (B,E). The modified Lhx9 isoforms were co-electroporated with the commissural axonal reporter eR3-123::mCherry. In control experiments (yellow arrows in B,E) all the axons cross the midline. commissural axons expressing the dominant negative isoforms turn also ipsilaterally (magenta arrows in A,D).

C. Quantification of the extent of commissural turning following ectopic expression of En-Lhx9 fusion protein. In control experiment 93.6+/-7.79% of En-Lhx9*m*-expressing commissural neurons cross the midline, while 55.1+/-21.3% of En-Lhx9-expressing neurons cross the midline.

F. Quantification of the extent of commissural turning following ectopic expression of Barhl2_En-Lhx9 fusion (BarLhx9) protein. In control experiment 99.5+/-0.8% of BarEn-Lhx9*m*-

expressing commissural neurons cross the midline, while 85.97+/-0.85% of BarEn-Lhx9-expressing neurons cross the midline.

Comparing the ratio of commissural axons between the controls (En-Lhx9*m* in C and BarEn-Lhx9*m* in F), and the experimental (En-Lhx9 in C and BarEn-Lhx9 in F), using Dunnett's method shows significant differences between the control and dominant negative Lhx9 isoforms. $P < 0.05$. (see Supplementary Statistics).

Fig. S11. Lhx1 activates expression downstream from the eR3-Lim enhancer.

The plasmids CAG::Lhx1-IRES-nEGFP (A,B), eR3-123::Cherry (A) and eR3-Lim (B) were electroporated into the chick spinal cord.

A. Ectopic expression of Lhx1 does not induce expression downstream of the eR3::123 enhancer.

B. Ectopic expression of Lhx1 induce expression downstream of the eR3::Lim enhancer.

**Supplementary Mathematical Analysis**

Analytical proof that negative feedback on Lhx2/9 restricts the width of the Robo3 pulse:

For the- iFFL without negative or positive feedback (system 2- iFFL):

Nullcline of L: $l' = p_1 - q_1 l = 0 \Rightarrow l = \frac{p_1}{q_1}$

Nullcline of R: $r' = p_3 \frac{l^{n_{lr}}}{l^{n_{lr}} + k^{lr^{n_{lr}}}} \frac{k^{br^{n_{br}}}}{b^{n_{br}} + k^{br^{n_{br}}}} - q_3 r = 0$

$$\Rightarrow r = \frac{p_3}{q_3} \frac{l^{n_{lr}}}{l^{n_{lr}} + k^{lr^{n_{lr}}}} \frac{k^{br^{n_{br}}}}{b^{n_{br}} + k^{br^{n_{br}}}}$$

$$\Rightarrow r \propto \frac{1}{b^{n_{br}}}$$

This shows that the rate of Robo3 repression decreases with Barhl1/2 expression. The system converges to a Robo3-off steady state after reaching a peak, with a decline rate $\frac{1}{b^{n_{br}}}$.

For the negative feedback enhanced iFFL (system 1- iFFL+dNFL):

Nullcline of L: $l' = p_1 \frac{k^{bl^{n_{bl}}}}{b^{n_{bl}} + k^{bl^{n_{bl}}}} - q_1 l = 0 \Rightarrow l = \frac{p_1}{q_1} \frac{k^{bl^{n_{bl}}}}{b^{n_{bl}} + k^{bl^{n_{bl}}}}$

$$\Rightarrow l \propto \frac{1}{b^{n_{bl}}}$$

Thus, $l \propto \frac{1}{b^{n_{bl}}}$ indicates that increased Barhl1/2 expression suppresses Lhx2/9 levels.

Nullcline of R: $r' = p_3 \frac{l^{n_{lr}}}{l^{n_{lr}} + k^{lr^{n_{lr}}}} \frac{k^{br^{n_{br}}}}{b^{n_{br}} + k^{br^{n_{br}}}} - q_3 r = 0$

$$\Rightarrow r = \frac{p_3}{q_3} \frac{l^{n_{lr}}}{l^{n_{lr}} + k^{lr^{n_{lr}}}} \frac{k^{br^{n_{br}}}}{b^{n_{br}} + k^{br^{n_{br}}}}$$

$$\Rightarrow r \propto \frac{1}{b^{n_{br}}}$$

In this NFL architecture, Robo3 again exhibits a transient pulse, but the decline occurs more rapidly due to the combined repressive effects on both Robo3 and its activator Lhx2/9. The effective decay rate is governed by the compound term $(\frac{1}{b^{n_{br}}}\frac{1}{b^{n_{bl}}})$, which is faster than the iFFL-only decay rate of $\frac{1}{b^{n_{br}}}$.

Therefore, although both architectures produce transient Robo3 responses, the presence of negative feedback in the NFL leads to a sharper and faster decline in Robo3 expression. This enhanced repression arises from earlier suppression of Lhx2/9, offering tighter control over pulse duration and preventing sustained activation of Robo3 as $\frac{1}{b^{n_{br}}}\frac{1}{b^{n_{bl}}} < \frac{1}{b^{n_{br}}}$.

#### Supplementary statistics

##### Fig. 1A

##### Quantiles

| Level | Minimum | 10% | 25% | Median | 75% | 90% | Maximum |
|---|---|---|---|---|---|---|---|
| E4.5 | 58.51614 | 66.74796 | 71.02055 | 75.7771 | 79.77528 | 82.65705 | 87.98283 |
| E5 | 50.54741 | 52.03787 | 56.2661 | 64.55393 | 70.52698 | 81.25333 | 86.78796 |
| E5.5 | 20.25046 | 36.18536 | 41.99179 | 49.56146 | 52.90674 | 59.05087 | 63.54715 |
| E6 | 18.74041 | 27.74686 | 32.13456 | 38.66621 | 46.28207 | 52.28833 | 53.64791 |
| E8 | 0.872541 | 1.363651 | 3.113553 | 5.339196 | 8.041627 | 13.51649 | 20.64777 |

##### Means and Std Deviations

| Level | Number | Mean | Std Dev | Std Err Mean | Lower 95% | Upper 95% |
|---|---|---|---|---|---|---|
| E4.5 | 55 | 74.827193 | 6.5605324 | 0.884622 | 73.053633 | 76.600754 |
| E5 | 41 | 64.654628 | 9.525967 | 1.4877061 | 61.647862 | 67.661394 |
| E5.5 | 56 | 48.025951 | 8.5531217 | 1.142959 | 45.735411 | 50.316492 |
| E6 | 67 | 38.978508 | 8.8713061 | 1.0838025 | 36.814627 | 41.14239 |
| E8 | 55 | 6.5752826 | 4.8174712 | 0.6495877 | 5.2729377 | 7.8776275 |

#### Fig. 2M

##### Quantiles

| Level | Minimum | 10% | 25% | Median | 75% | 90% | Maximum |
|---|---|---|---|---|---|---|---|
| Lhx2 | 1.06383 | 1.300629 | 1.551423 | 1.835034 | 2.162532 | 2.84737 | 3.458333 |
| Lhx9 | 0.833333 | 0.974089 | 1.195122 | 1.96875 | 2.527273 | 2.927644 | 3.941176 |
| nGFP | 0.686047 | 0.842105 | 0.925373 | 0.975309 | 1.057143 | 1.244186 | 1.509804 |

##### Means and Std Deviations

| Level | Number | Mean | Std Dev | Std Err Mean | Lower 95% | Upper 95% |
|---|---|---|---|---|---|---|
| Lhx2 | 88 | 1.9207057 | 0.5276952 | 0.0562525 | 1.8088977 | 2.0325136 |
| Lhx9 | 55 | 1.941702 | 0.8031914 | 0.1083023 | 1.7245689 | 2.1588351 |
| nGFP | 79 | 1.0055976 | 0.1479301 | 0.0166434 | 0.972463 | 1.0387321 |

##### Means Comparisons
##### Comparisons with a control using Dunnett's Method

Control Group = nGFP

##### Confidence Quantile

| \|d\| | Alpha |
|---|---|
| 2.23028 | 0.05 |

##### LSD Threshold Matrix

| Level | Abs(Dif)-LSD | p-Value |
|---|---|---|
| Lhx9 | 0.73 | <.0001* |
| Lhx2 | 0.733 | <.0001* |
| nGFP | -0.19 | 1.0000 |

Positive values show pairs of means that are significantly different.

#### Fig. 2N

##### Quantiles

| Level | Minimum | 10% | 25% | Median | 75% | 90% | Maximum |
|---|---|---|---|---|---|---|---|
| Lhx2 | 0.974359 | 1.773913 | 2.309495 | 2.534884 | 3.228523 | 3.550831 | 3.679245 |
| Lhx9 | 1.133333 | 1.286229 | 1.528433 | 2.232303 | 3.153928 | 3.782095 | 4.153846 |
| nGFP | 0.719298 | 0.799564 | 0.925923 | 0.99375 | 1.052807 | 1.216279 | 1.34 |

##### Means and Std Deviations

| Level | Number | Mean | Std Dev | Std Err Mean | Lower 95% | Upper 95% |
|---|---|---|---|---|---|---|
| Lhx2 | 21 | 2.6252672 | 0.6672333 | 0.1456023 | 2.3215463 | 2.9289882 |
| Lhx9 | 28 | 2.3856999 | 0.8889014 | 0.1679866 | 2.04102 | 2.7303799 |
| nGFP | 34 | 0.9989007 | 0.138212 | 0.0237032 | 0.9506762 | 1.0471251 |

##### Means Comparisons
##### Comparisons with a control using Dunnett's Method

Control Group = nGFP

##### Confidence Quantile

| \|d\| | Alpha |
|---|---|
| 2.26071 | 0.05 |

##### LSD Threshold Matrix

| Level | Abs(Dif)-LSD | p-Value |
|---|---|---|
| Lhx2 | 1.237 | <.0001* |
| Lhx9 | 1.028 | <.0001* |
| nGFP | -0.34 | 1.0000 |

Positive values show pairs of means that are significantly different.

Fig. 2O

#### Quantiles

| Level | Minimum | 10% | 25% | Median | 75% | 90% | Maximum |
|---|---|---|---|---|---|---|---|
| Barhl1 | 0.083333 | 0.315302 | 0.446769 | 0.562873 | 0.636343 | 0.720765 | 0.886364 |
| Barhl2 | 0.029412 | 0.090505 | 0.163217 | 0.307692 | 0.431538 | 0.673148 | 1.263158 |
| nGFP | 0.615385 | 0.88254 | 0.952381 | 1 | 1.0625 | 1.150857 | 1.333333 |

#### Means and Std Deviations

| Level | Number | Mean | Std Dev | Std Err Mean | Lower 95% | Upper 95% |
|---|---|---|---|---|---|---|
| Barhl1 | 44 | 0.5428589 | 0.158907 | 0.0239561 | 0.4945468 | 0.5911711 |
| Barhl2 | 77 | 0.3464894 | 0.2515665 | 0.0286687 | 0.2893908 | 0.403588 |
| nGFP | 51 | 1.000881 | 0.1102084 | 0.0154323 | 0.9698844 | 1.0318777 |

#### Means Comparisons

#### Comparisons with a control using Dunnett's Method

Control Group = nGFP

#### Confidence Quantile

| \|d\| | Alpha |
|---|---|
| 2.22733 | 0.05 |

#### LSD Threshold Matrix

| Level | Abs(Dif)-LSD | p-Value |
|---|---|---|
| nGFP | -0.09 | 1.0000 |
| Barhl1 | 0.368 | <.0001* |
| Barhl2 | 0.576 | <.0001* |

Positive values show pairs of means that are significantly different.

Fig. 2P

##### Quantiles

| Level | Minimum | 10% | 25% | Median | 75% | 90% | Maximum |
|---|---|---|---|---|---|---|---|
| Barhl1 | 1.089286 | 1.236786 | 1.338636 | 1.67803 | 2.175245 | 2.796875 | 3.964286 |
| nGFP | 0.719298 | 0.799564 | 0.925923 | 0.99375 | 1.052807 | 1.216279 | 1.34 |

##### Means and Std Deviations

| Level | Number | Mean | Std Dev | Std Err Mean | Lower 95% | Upper 95% |
|---|---|---|---|---|---|---|
| Barhl1 | 28 | 1.8717637 | 0.6942424 | 0.1311995 | 1.6025646 | 2.1409627 |
| nGFP | 34 | 0.9989007 | 0.138212 | 0.0237032 | 0.9506762 | 1.0471251 |

##### Means Comparisons
##### Comparisons with a control using Dunnett's Method

Control Group = nGFP

##### Confidence Quantile

| \|d\| | Alpha |
|---|---|
| 2.00030 | 0.05 |

##### LSD Threshold Matrix

| Level | Abs(Dif)-LSD | p-Value |
|---|---|---|
| Barhl1 | 0.629 | <.0001* |
| nGFP | -0.23 | 1.0000 |

Positive values show pairs of means that are significantly different.

Fig. 2Q

#### Quantiles

| Level | Minimum | 10% | 25% | Median | 75% | 90% | Maximum |
|---|---|---|---|---|---|---|---|
| Barhl2 | 1.613861 | 1.822857 | 2.181939 | 2.491071 | 3.048665 | 3.902012 | 4.020619 |
| nGFP | 0.686047 | 0.842105 | 0.925373 | 0.975309 | 1.057143 | 1.244186 | 1.509804 |

#### Means and Std Deviations

| Level | Number | Mean | Std Dev | Std Err Mean | Lower 95% | Upper 95% |
|---|---|---|---|---|---|---|
| Barhl2 | 30 | 2.6478952 | 0.6871437 | 0.1254547 | 2.3913116 | 2.9044789 |
| nGFP | 79 | 1.0055976 | 0.1479301 | 0.0166434 | 0.972463 | 1.0387321 |

#### Means Comparisons

#### Comparisons with a control using Dunnett's Method

Control Group = nGFP

#### Confidence Quantile

| \|d\| | Alpha |
|---|---|
| 1.98238 | 0.05 |

#### LSD Threshold Matrix

| Level | Abs(Dif)-LSD | p-Value |
|---|---|---|
| Barhl2 | 1.481 | <.0001* |
| nGFP | -0.12 | 1.0000 |

Positive values show pairs of means that are significantly different.

Fig. 3D

**Quantiles**

| Level | Minimum | 10% | 25% | Median | 75% | 90% | Maximum |
|---|---|---|---|---|---|---|---|
| dI1-Barhl2 | 0.004421 | 0.023627 | 0.047376 | 0.10618 | 0.14533 | 0.187178 | 0.266697 |
| dI1-Barhl2+Robo3 | 0.222222 | 0.458058 | 0.557271 | 0.705444 | 0.798477 | 0.888489 | 0.999107 |
| dI1-cherry | 0.173158 | 0.245613 | 0.329382 | 0.5 | 0.618222 | 0.73955 | 0.856465 |

**Means and Std Deviations**

| Level | Number | Mean | Std Dev | Std Err Mean | Lower 95% | Upper 95% |
|---|---|---|---|---|---|---|
| dI1-Barhl2 | 66 | 0.1034786 | 0.0620562 | 0.0076386 | 0.0882233 | 0.1187339 |
| dI1-Barhl2+Robo3 | 118 | 0.6835434 | 0.1749346 | 0.016104 | 0.6516502 | 0.7154366 |
| dI1-cherry | 54 | 0.4880303 | 0.1875974 | 0.0255288 | 0.436826 | 0.5392345 |

**LSD Threshold Matrix**

| Level | Abs(Dif)-LSD | p-Value |
|---|---|---|
| dI1-Barhl2+Robo3 | 0.139 | <.0001* |
| dI1-cherry | -0.07 | 1.0000 |
| dI1-Barhl2 | 0.321 | <.0001* |

Positive values show pairs of means that are significantly different

Fig. 3H

**Quantiles**

| Level | Minimum | 10% | 25% | Median | 75% | 90% | Maximum |
|---|---|---|---|---|---|---|---|
| Barhl2 | 0.094104 | 0.101579 | 0.126039 | 0.214947 | 0.25421 | 0.34504 | 0.379531 |
| GFP | 0.343795 | 0.443783 | 0.520448 | 0.569479 | 0.644966 | 0.66384 | 0.747904 |

**Means and Std Deviations**

| Level | Number | Mean | Std Dev | Std Err Mean | Lower 95% | Upper 95% |
|---|---|---|---|---|---|---|
| Barhl2 | 24 | 0.2078383 | 0.0823777 | 0.0168153 | 0.1730532 | 0.2426234 |
| GFP | 19 | 0.5718118 | 0.0979548 | 0.0224724 | 0.524599 | 0.6190245 |

**LSD Threshold Matrix**

| Level | Abs(Dif)-LSD | p-Value |
|---|---|---|
| GFP | -0.06 | 1.0000 |
| Barhl2 | 0.308 | <.0001* |

Positive values show pairs of means that are significantly different.

Fig. 3K

**Quantiles**

| Level | Minimum | 10% | 25% | Median | 75% | 90% | Maximum |
|---|---|---|---|---|---|---|---|
| dI2-Barhl2 | 0 | 0.002132 | 0.013996 | 0.085647 | 0.222622 | 0.276671 | 0.410784 |
| dI2-GFP | 0.490279 | 0.544003 | 0.632414 | 0.723577 | 0.817862 | 0.847518 | 0.956213 |

#### Means and Std Deviations

| Level | Number | Mean | Std Dev | Std Err Mean | Lower 95% | Upper 95% |
|---|---|---|---|---|---|---|
| dI2-Barhl2 | 25 | 0.1171041 | 0.1176354 | 0.0235271 | 0.0685466 | 0.1656616 |
| dI2-GFP | 48 | 0.7113492 | 0.1121145 | 0.0161823 | 0.6787945 | 0.7439038 |

#### LSD Threshold Matrix

| Level | Abs(Dif)-LSD | p-Value |
|---|---|---|
| dI2-GFP | -0.05 | 1.0000 |
| dI2-Barhl2 | 0.538 | <.0001* |

Positive values show pairs of means that are significantly different

Fig. 3M

**Quantiles**

| Level | Minimum | 10% | 25% | Median | 75% | 90% | Maximum |
|---|---|---|---|---|---|---|---|
| dI1-Barhl2 | 0 | 0 | 0.006865 | 0.022317 | 0.054575 | 0.125648 | 0.321183 |
| dI1-Barhl2DEng | 0.238451 | 0.373043 | 0.482894 | 0.615124 | 0.730357 | 0.875585 | 0.960252 |
| dI1-cherry | 0.173158 | 0.243548 | 0.301044 | 0.485263 | 0.591061 | 0.695419 | 0.748548 |

**Means and Std Deviations**

| Level | Number | Mean | Std Dev | Std Err Mean | Lower 95% | Upper 95% |
|---|---|---|---|---|---|---|
| dI1-Barhl2 | 95 | 0.0458205 | 0.0671411 | 0.0068885 | 0.0321432 | 0.0594978 |
| dI1-Barhl2DEng | 85 | 0.6072762 | 0.1762757 | 0.0191198 | 0.5692544 | 0.645298 |
| dI1-cherry | 50 | 0.4595087 | 0.1638733 | 0.0231752 | 0.4129365 | 0.506081 |

**LSD Threshold Matrix**

| Level | Abs(Dif)-LSD | p-Value |
|---|---|---|
| dI1-Barhl2DEng | 0.087 | <.0001* |
| dI1-cherry | -0.07 | 1.0000 |
| dI1-Barhl2 | 0.354 | <.0001* |

Positive values show pairs of means that are significantly different.

Fig. 4H

#### Quantiles

| Level | Minimum | 10% | 25% | Median | 75% | 90% | Maximum |
|---|---|---|---|---|---|---|---|
| GFP | 0.091128 | 0.105096 | 0.161224 | 0.295526 | 0.424068 | 0.507742 | 0.576465 |
| Lhx2/9 | 0.141703 | 0.159546 | 0.20456 | 0.235459 | 0.284274 | 0.366461 | 0.38843 |

#### Means and Std Deviations

| Level | Number | Mean | Std Dev | Std Err Mean | Lower 95% | Upper 95% |
|---|---|---|---|---|---|---|
| GFP | 49 | 0.3022422 | 0.1466573 | 0.020951 | 0.2601173 | 0.3443671 |
| Lhx2/9. | 27 | 0.2514383 | 0.0674453 | 0.0129799 | 0.2247578 | 0.2781188 |

##### Means Comparisons
##### Comparisons with a control using Dunnett's Method

Control Group = GFP

| Level | Abs(Dif)-LSD | p-Value |
|---|---|---|
| GFP | -0.05 | 1 |
| Lhx2/9. | -0.01 | 0.0924 |

Fig. 6D

**Quantiles**

| Level | Minimum | 10% | 25% | Median | 75% | 90% | Maximum |
|---|---|---|---|---|---|---|---|
| eR3+Lhx9 | 0.17473 | 0.309493 | 0.431159 | 0.604749 | 0.72252 | 0.748999 | 0.853113 |
| eR3+Lhx9m | 0.885498 | 0.913046 | 0.973006 | 0.992757 | 1 | 1 | 1 |
| eR3-LimM | 0.081487 | 0.151458 | 0.239518 | 0.414478 | 0.564283 | 0.684303 | 0.770237 |

**Means and Std Deviations**

| Level | Number | Mean | Std Dev | Std Err Mean | Lower 95% | Upper 95% |
|---|---|---|---|---|---|---|
| eR3+Lhx9 | 20 | 0.566004 | 0.1769903 | 0.0395762 | 0.48317 | 0.648838 |
| eR3+Lhx9m | 43 | 0.9784779 | 0.032161 | 0.0049045 | 0.9685802 | 0.9883756 |
| eR3-LimM | 44 | 0.4104154 | 0.1918995 | 0.0289299 | 0.3520726 | 0.4687582 |

**LSD Threshold Matrix**

| Level | Abs(Dif)-LSD | p-Value |
|---|---|---|
| eR3+Lhx9m | -0.07 | 1 |
| eR3+Lhx9 | 0.323 | <.0001* |
| eR3-LimM | 0.497 | <.0001* |

Positive values show pairs of means that are significantly different

Fig. 6E

| DNA | Lhx9 | N sections | Mean(& fraction) | Median(& fraction) |
|---|---|---|---|---|
| 1 | 9 | 7 | 0.044964779 | 0.05 |
| 1 | 9m | 7 | 0.018669671 | 0.017848029 |
| 2 | 9 | 7 | 0.055302518 | 0.052456324 |
| 2 | 9m | 7 | 0.01467405 | 0.016491247 |
| 3 | 9 | 7 | 0.102344686 | 0.093563479 |
| 3 | 9m | 7 | 0.027645931 | 0.021710109 |
| 4 | 9 | 7 | 0.11336167 | 0.114682679 |
| 4 | 9m | 7 | 0.024262359 | 0.021500464 |
| 5 | 9 | 7 | 0.103632743 | 0.112 |
| 5 | 9m | 7 | 0.037908832 | 0.01727556 |
| 6 | 9 | 6 | 0.135684572 | 0.147684885 |
| 6 | 9m | 7 | 0.028451862 | 0.025517079 |
| 7 | 9 | 7 | 0.056691486 | 0.053061485 |
| 7 | 9m | 7 | 0.025041582 | 0.021648998 |
| 8 | 9 | 7 | 0.034468575 | 0.032762572 |
| 8 | 9m | 7 | 0.022051246 | 0.017881983 |
| 9 | 9 | 7 | 0.049285256 | 0.05 |
| 9 | 9m | 7 | 0.019132471 | 0.01661585 |
| 10 | 9 | 3 | 0.018043689 | 0.018831899 |
| 10 | 9m | 3 | 0.013739105 | 0.013358738 |

Fig. S4E

| Level | Number | Mean | Std Dev | Std Err Mean | Lower 95% | Upper 95% |
|---|---|---|---|---|---|---|
| GFP | 19 | 60.7341 | 11.2375 | 2.5781 | 66.15 | 55.318 |
| Lhx2. | 15 | 46.9731 | 12.1055 | 3.1256 | 53.677 | 40.269 |
| Lhx2/9. | 33 | 40.9586 | 15.6385 | 2.7223 | 46.504 | 35.413 |
| Lhx9. | 34 | 45.8174 | 10.9004 | 1.8694 | 49.621 | 42.014 |

Fig. S8D

#### Means and Std Dev

| Embryo | % comm | N sections | Mean(Data) | Std Dev(Data) |
|---|---|---|---|---|
| A | CAG::EGFP | 5 | 0.564159183 | 0.126944893 |
| A | eR3-Lim::mCherry | 5 | 0.746453587 | 0.180986647 |
| B | CAG::EGFP | 6 | 0.776866676 | 0.15116649 |
| B | eR3-Lim::mCherry | 6 | 0.912926373 | 0.113938276 |
| C | CAG::EGFP | 10 | 0.6858908 | 0.092421571 |
| C | eR3-Lim::mCherry | 10 | 0.854426333 | 0.116535125 |
| D | CAG::EGFP | 8 | 0.5247755 | 0.122656685 |
| D | eR3-Lim::mCherry | 8 | 0.758714091 | 0.203488211 |
| E | CAG::EGFP | 8 | 0.717708055 | 0.077852529 |
| E | eR3-Lim::mCherry | 8 | 0.952061352 | 0.04103569 |
| F | CAG::EGFP | 4 | 0.677565491 | 0.108219772 |
| F | eR3-Lim::mCherry | 4 | 0.945694378 | 0.060560937 |

#### Variability Chart for Data

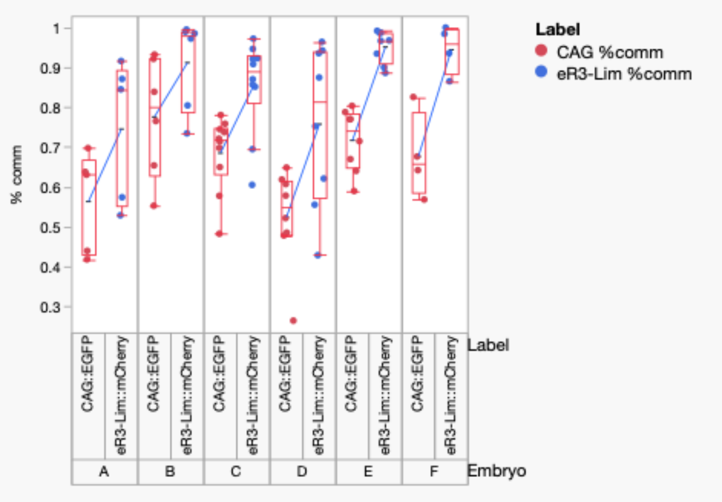

Fig. S10C

**Quantiles**

| Level | Minimum | 10% | 25% | Median | 75% | 90% | Maximum |
|---|---|---|---|---|---|---|---|
| EnLhx9 | 0.081614 | 0.244399 | 0.457303 | 0.53848 | 0.640616 | 0.898425 | 0.966242 |
| EnLhx9m | 0.71515 | 0.787738 | 0.893938 | 0.983824 | 0.99356 | 0.999714 | 0.999922 |

**Means and Std Deviations**

| Level | Number | Mean | Std Dev | Std Err Mean | Lower 95% | Upper 95% |
|---|---|---|---|---|---|---|
| EnLhx9 | 26 | 0.5508412 | 0.2131236 | 0.041797 | 0.4647587 | 0.6369237 |
| EnLhx9m | 33 | 0.9368652 | 0.0779769 | 0.013574 | 0.9092158 | 0.9645146 |

#### LSD Threshold Matrix

| Level | Abs(Dif)-LSD | p-Value |
|---|---|---|
| EnLhx9m | -0.08 | 1 |
| EnLhx9 | 0.306 | <.0001* |

Positive values show pairs of means that are significantly different

Fig. S10F

**Quantiles**

| Level | Minimum | 10% | 25% | Median | 75% | 90% | Maximum |
|---|---|---|---|---|---|---|---|
| BarEn-Lhx9 | 0.693187 | 0.728499 | 0.803502 | 0.865781 | 0.925768 | 0.964482 | 0.991222 |
| BarEn-Lhx9m | 0.965322 | 0.981202 | 0.995958 | 0.998059 | 0.999975 | 1 | 1 |

**Means and Std Deviations**

| **Level** | **Number** | **Mean** | **Std Dev** | **Std Err Mean** | **Lower 95%** | **Upper 95%** |
|---|---|---|---|---|---|---|
| BarEn-Lhx9 | 36 | 0.8597496 | 0.0821162 | 0.013686 | 0.8319654 | 0.8875337 |
| BarEn-Lhx9m | 25 | 0.994907 | 0.0085735 | 0.0017147 | 0.991368 | 0.998446 |

**LSD Threshold Matrix**

| Level | Abs(Dif)-LSD | p-Value |
|---|---|---|
| BarEn-Lhx9m | -0.04 | 1 |
| BarEn-Lhx9 | 0.102 | <.0001* |

Supp. Fig. S1

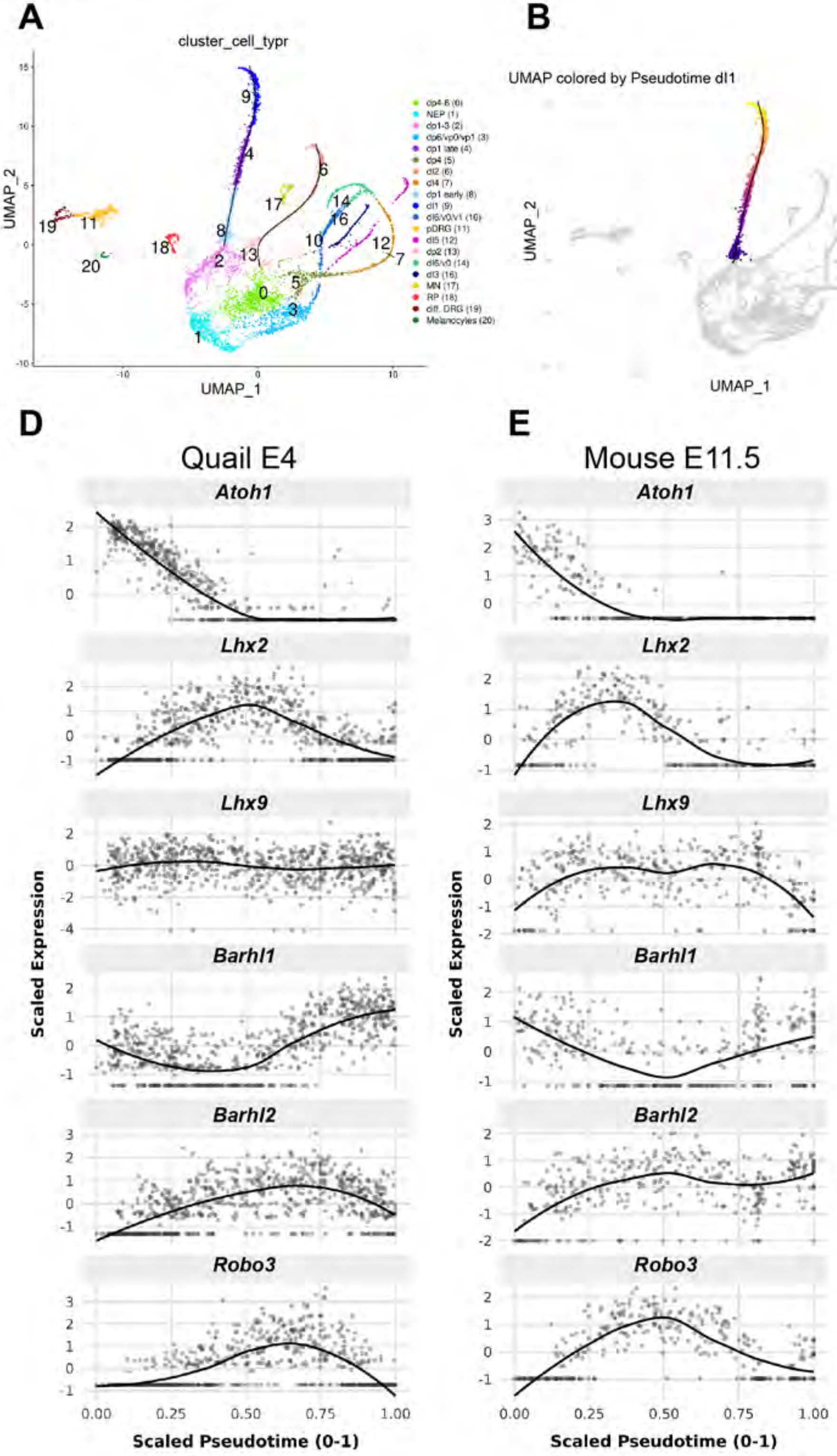
A
cluster_cell_typr
UMAP_2
UMAP_1
dp4-6 (0)
NEP (1)
dp1-3 (2)
dp6/vp0/vp1 (3)
dp1 late (4)
dp4 (5)
dI2 (6)
dI4 (7)
dp1 early (8)
dI1 (9)
dI6/V0/V1 (10)
pDRG (11)
dI5 (12)
dp2 (13)
dI6/v0 (14)
dI3 (16)
MN (17)
RP (18)
diff. DRG (19)
Melanocytes (20)
B
UMAP colored by Pseudotime dI1
C
UMAP colored by Pseudotime dI2
D
Quail E4
E
Mouse E11.5
Atoh1
Lhx2
Lhx9
Barhl1
Barhl2
Robo3
Scaled Expression
Scaled Pseudotime (0-1)

Supp Fig. S2

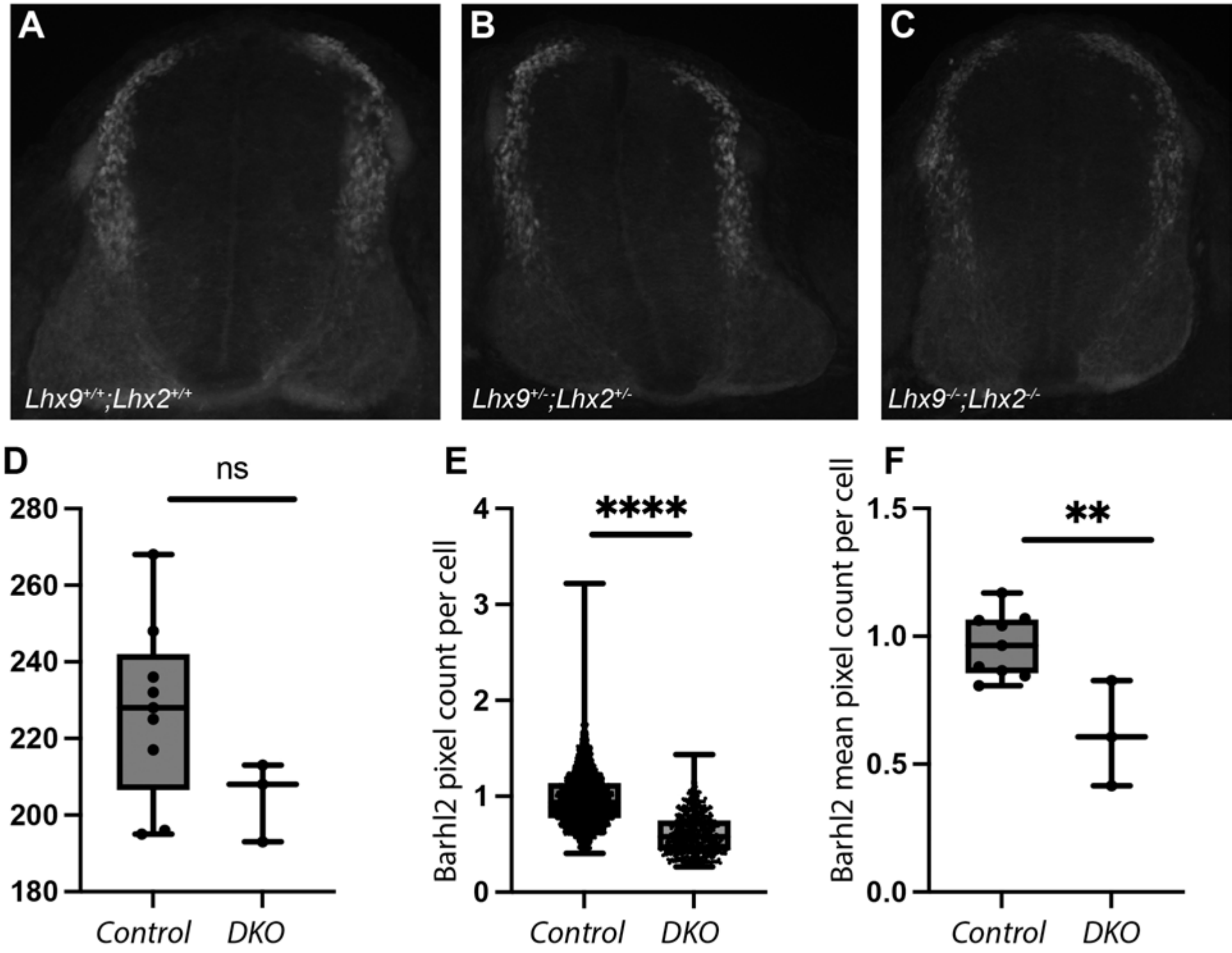
A
B
C
Lhx9+/+;Lhx2+/+
Lhx9+/-;Lhx2+/-
Lhx9-/-;Lhx2-/-
D
ns
### of Barhl2 cells
280
260
240
220
200
180
Control
DKO
E
****
Barhl2 pixel count per cell
4
3
2
1
0
Control
DKO
F
**
Barhl2 mean pixel count per cell
1.5
1.0
0.5
0.0
Control
DKO

Supp Fig. S3

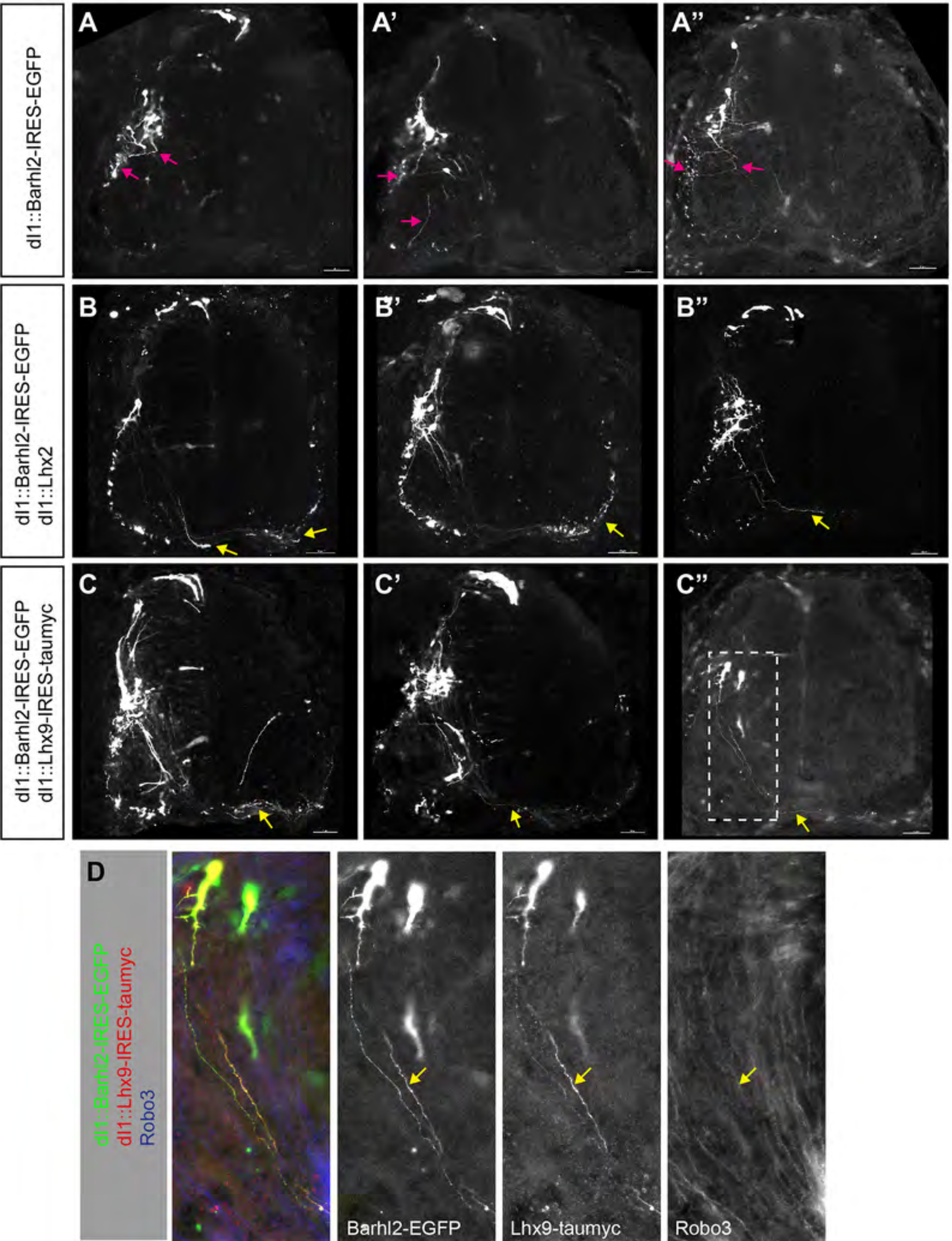
dI1::Barhl2-IRES-EGFP
A
A'
A''
dI1::Barhl2-IRES-EGFP
dI1::Lhx2
B
B'
B''
dI1::Barhl2-IRES-EGFP
dI1::Lhx9-IRES-taumyc
C
C'
C''
D
dI1::Barhl2-IRES-EGFP
dI1::Lhx9-IRES-taumyc
Robo3
Barhl2-EGFP
Lhx9-taumyc
Robo3

Supp Fig. S4

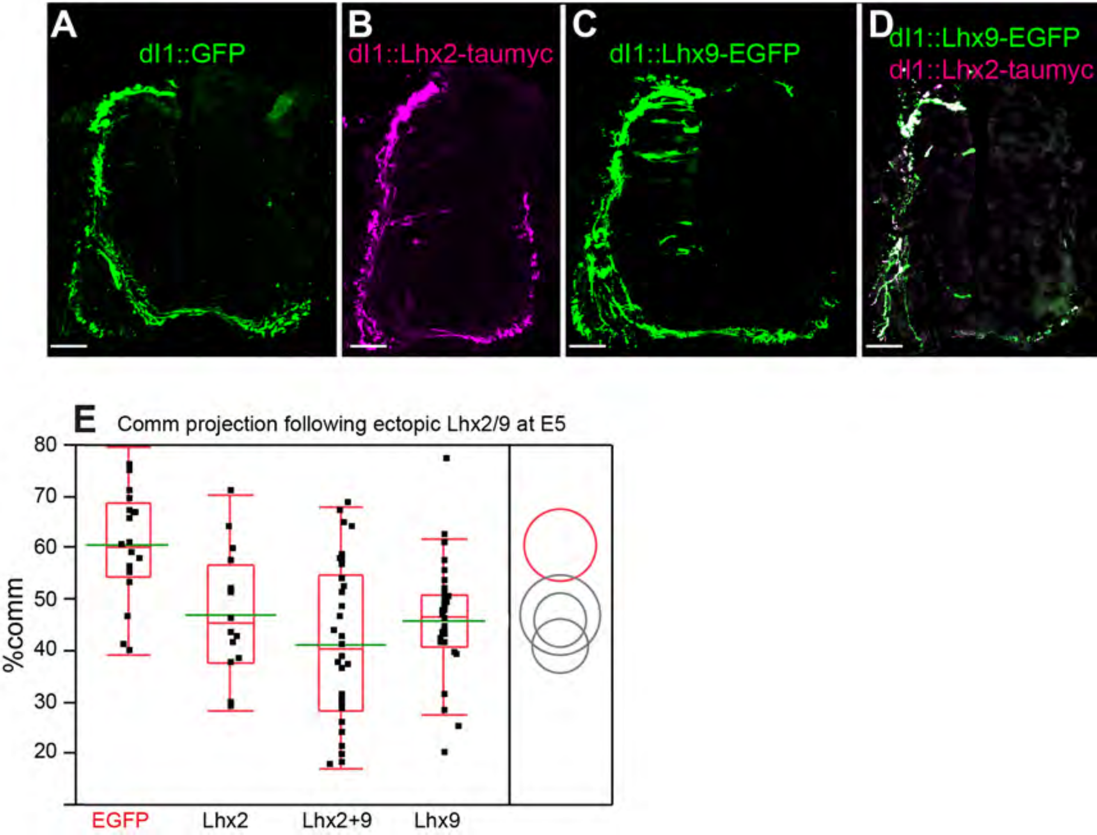
A
dI1::GFP
B
dI1::Lhx2-taumyc
C
dI1::Lhx9-EGFP
D
dI1::Lhx9-EGFP
dI1::Lhx2-taumyc
E
Comm projection following ectopic Lhx2/9 at E5
%comm
80
70
60
50
40
30
20
EGFP
Lhx2
Lhx2+9
Lhx9

Supp Fig. S5

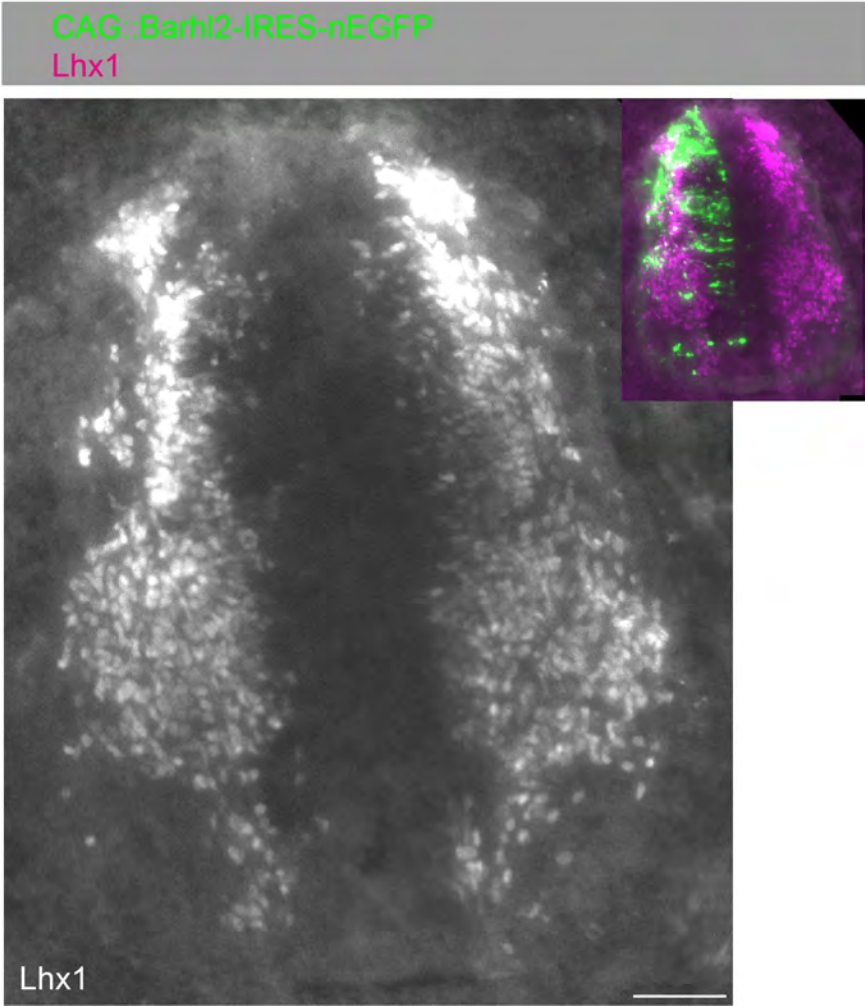

### Fig. S6

**A**

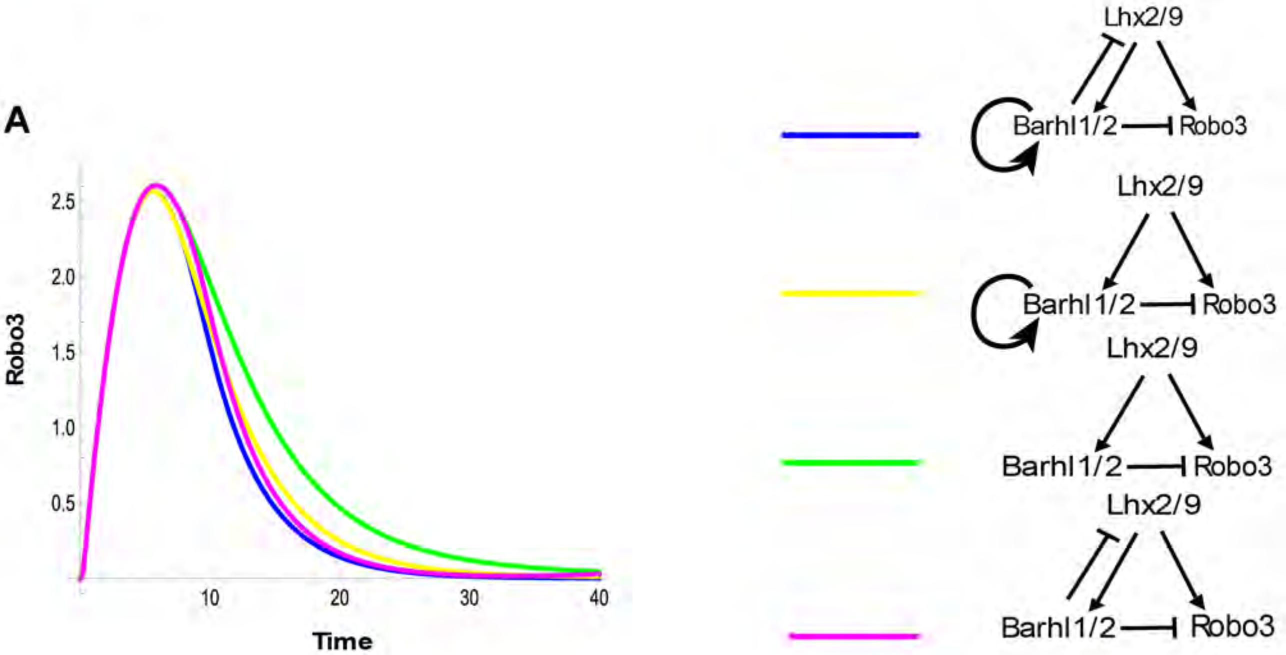
Robo3
Time
2.5
2.0
1.5
1.0
0.5
10
20
30
40
Lhx2/9
Barhl1/2
Robo3
Lhx2/9
Barhl1/2
Robo3
Lhx2/9
Barhl1/2
Robo3
Lhx2/9
Barhl1/2
Robo3

**B**

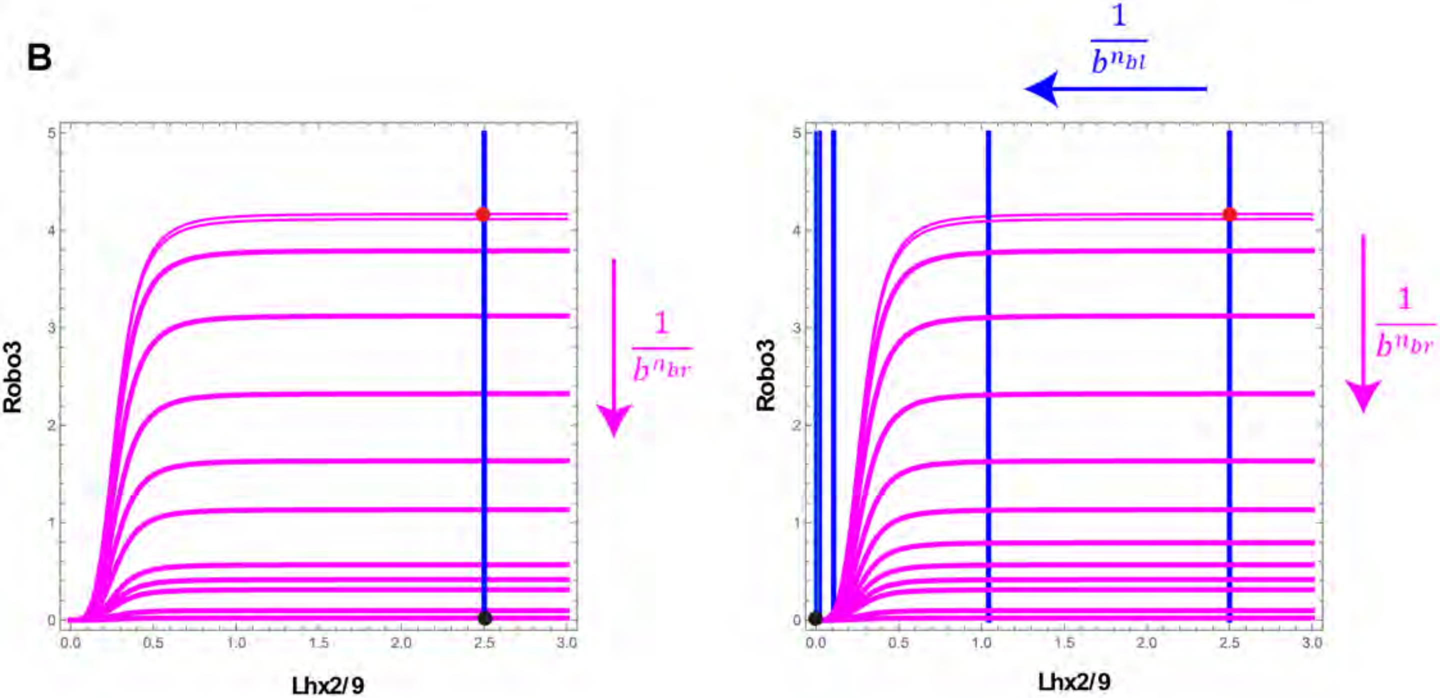
Robo3
Lhx2/9
$\frac{1}{b^{n_{br}}}$
$\frac{1}{b^{n_{bl}}}$
Robo3
Lhx2/9
$\frac{1}{b^{n_{br}}}$

Supp Fig. S7

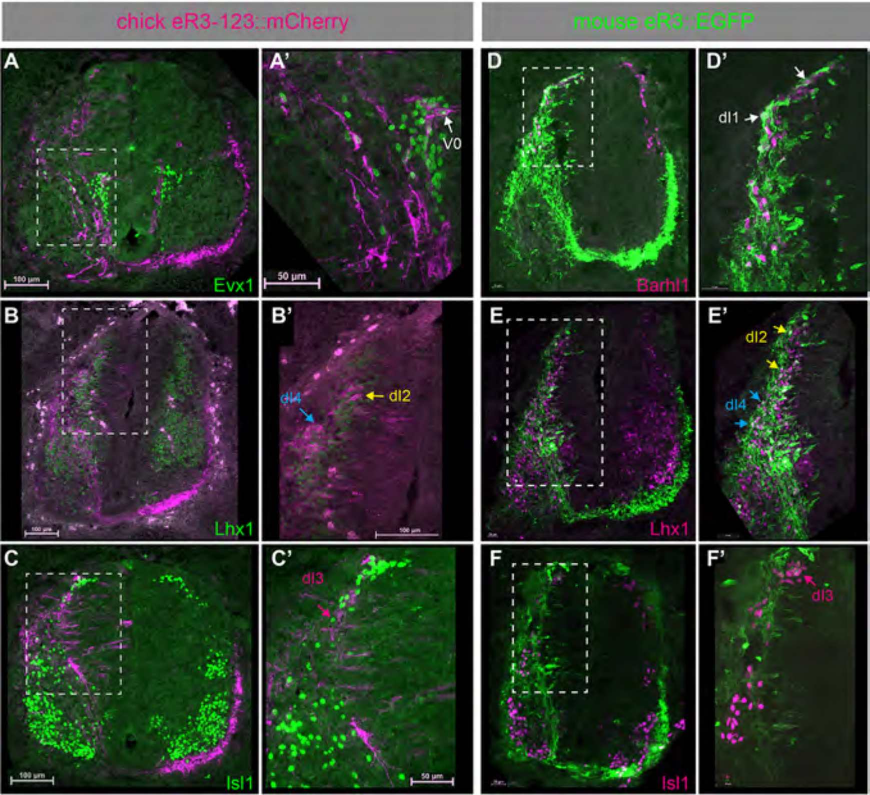
chick eR3-123::mCherry
mouse eR3::EGFP
A
A'
V0
100 µm
Evx1
50 µm
D
D'
dI1
Barhl1
B
B'
dI4
dI2
100 µm
Lhx1
E
E'
dI2
dI4
Lhx1
C
C'
dI3
100 µm
Isl1
50 µm
F
F'
dI3
Isl1

Supp Fig. S8

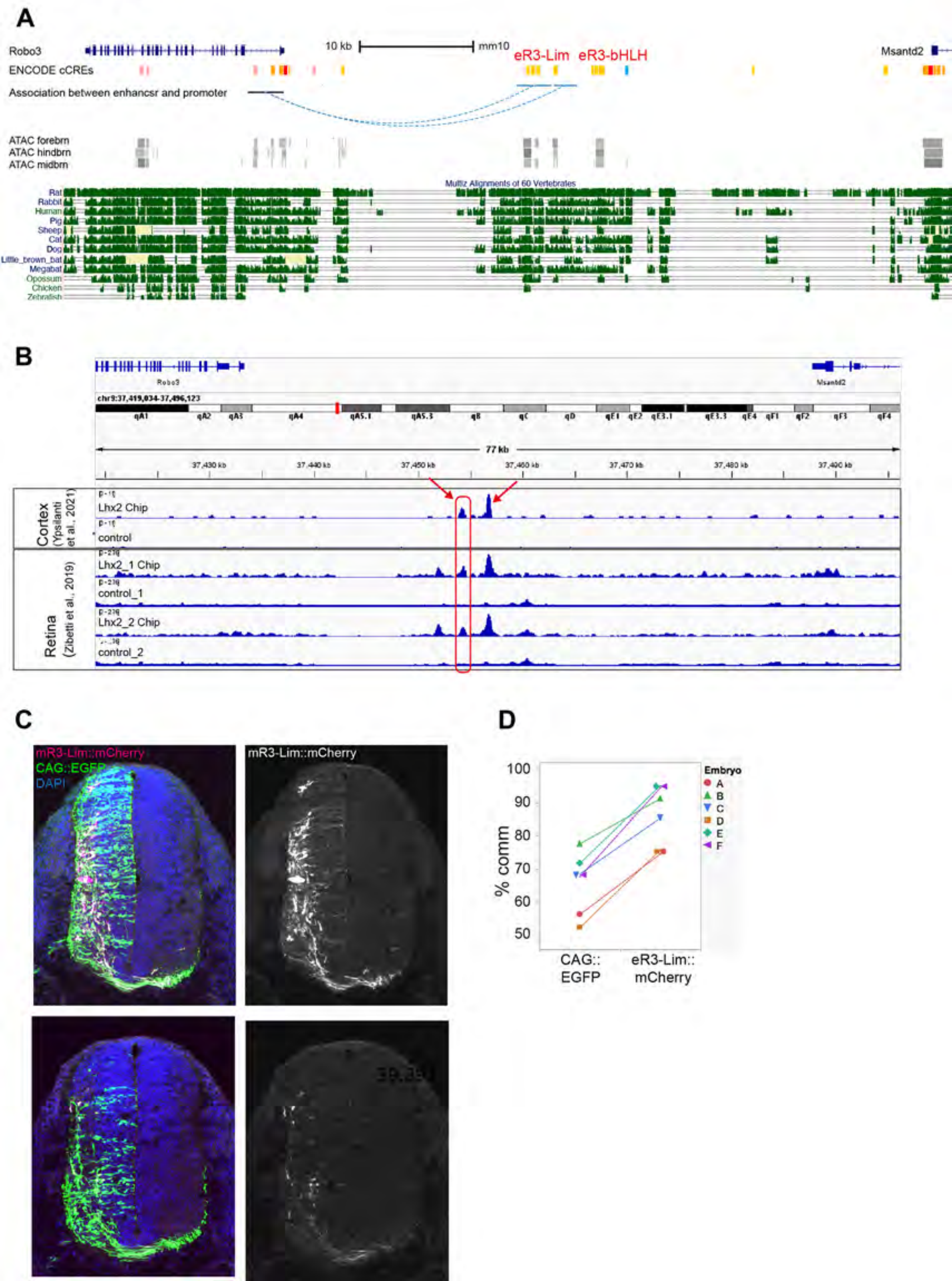

A
Robo3
10 kb
mm10
eR3-Lim
eR3-bHLH
Msantd2
ENCODE cCREs
Association between enhancsr and promoter
ATAC forebrn
ATAC hindbrn
ATAC midbrn
Multiz Alignments of 60 Vertebrates
Rat
Rabbit
Human
Pig
Sheep
Cat
Dog
Little_brown_bat
Megabat
Opossum
Chicken
Zebrafish
B
Robo3
Msantd2
chr9:37,419,034-37,496,123
77 kb
37,430 kb
37,440 kb
37,450 kb
37,460 kb
37,470 kb
37,480 kb
Cortex (Ypsilanti et al., 2021)
Lhx2 Chip
control
Retina (Zibetti et al., 2019)
Lhx2_1 Chip
control_1
Lhx2_2 Chip
control_2
C
mR3-Lim::mCherry
CAG::EGFP
DAPI
mR3-Lim::mCherry
D
100
90
80
70
60
50
% comm
CAG:: EGFP
eR3-Lim:: mCherry
Embryo
A
B
C
D
E
F

#### Supp Fig. S9

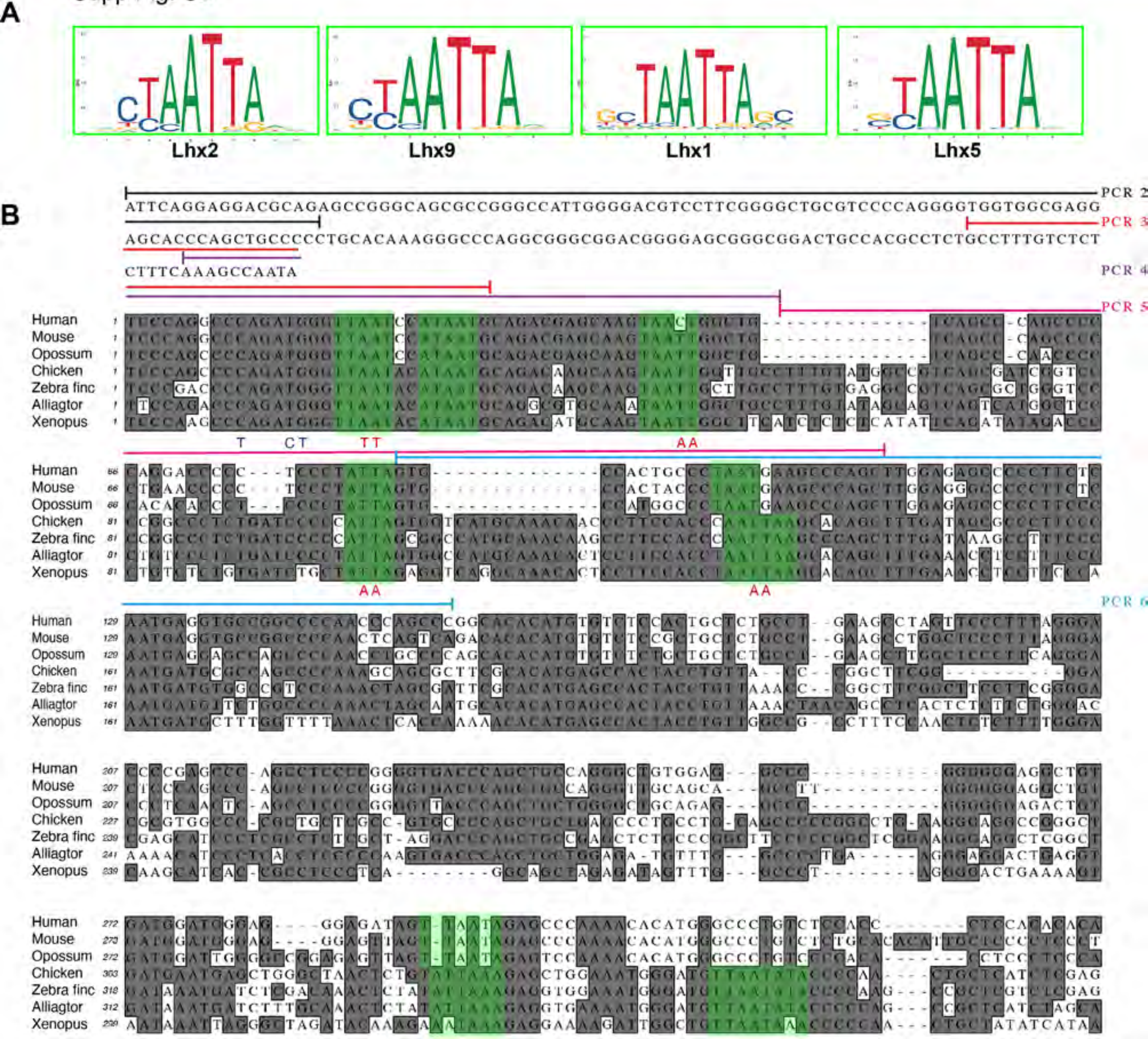
A
Lhx2
Lhx9
Lhx1
Lhx5
B
PCR 2
ATTCAGGAGGACGCAGAGCCGGGCAGCGCCGGGCCATTGGGGACGTCCTTCGGGGCTGCGTCCCCAGGGGTGGTGGCGAGG
PCR 3
AGCACCCAGCTGCCCCCTGCACAAAGGGCCCAGGCGGGCGGACGGGGAGCGGGCGGACTGCCACGCCTCTGCCTTTGTCTCT
CTTTCAAAGCCAATA
PCR 4
PCR 5
PCR 6
Human
Mouse
Opossum
Chicken
Zebra finc
Alliagtor
Xenopus

Supp Fig. S10.

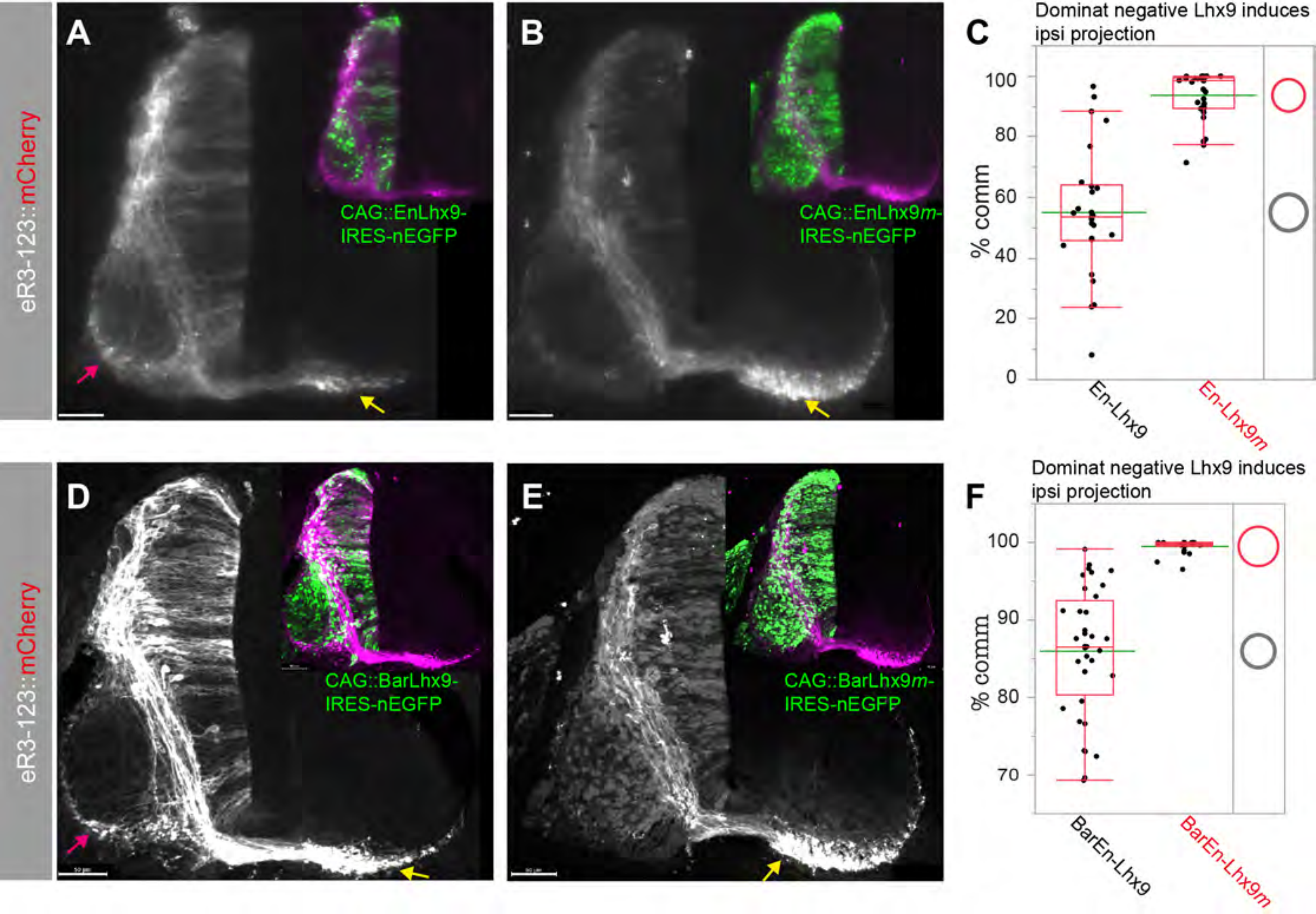
eR3-123::mCherry
A
CAG::EnLhx9-
IRES-nEGFP
B
CAG::EnLhx9m-
IRES-nEGFP
C
Dominat negative Lhx9 induces
ipsi projection
% comm
100
80
60
40
20
0
En-Lhx9
En-Lhx9m
eR3-123::mCherry
D
CAG::BarLhx9-
IRES-nEGFP
E
CAG::BarLhx9m-
IRES-nEGFP
F
Dominat negative Lhx9 induces
ipsi projection
% comm
100
90
80
70
BarEn-Lhx9
BarEn-Lhx9m

Supp Fig. S11

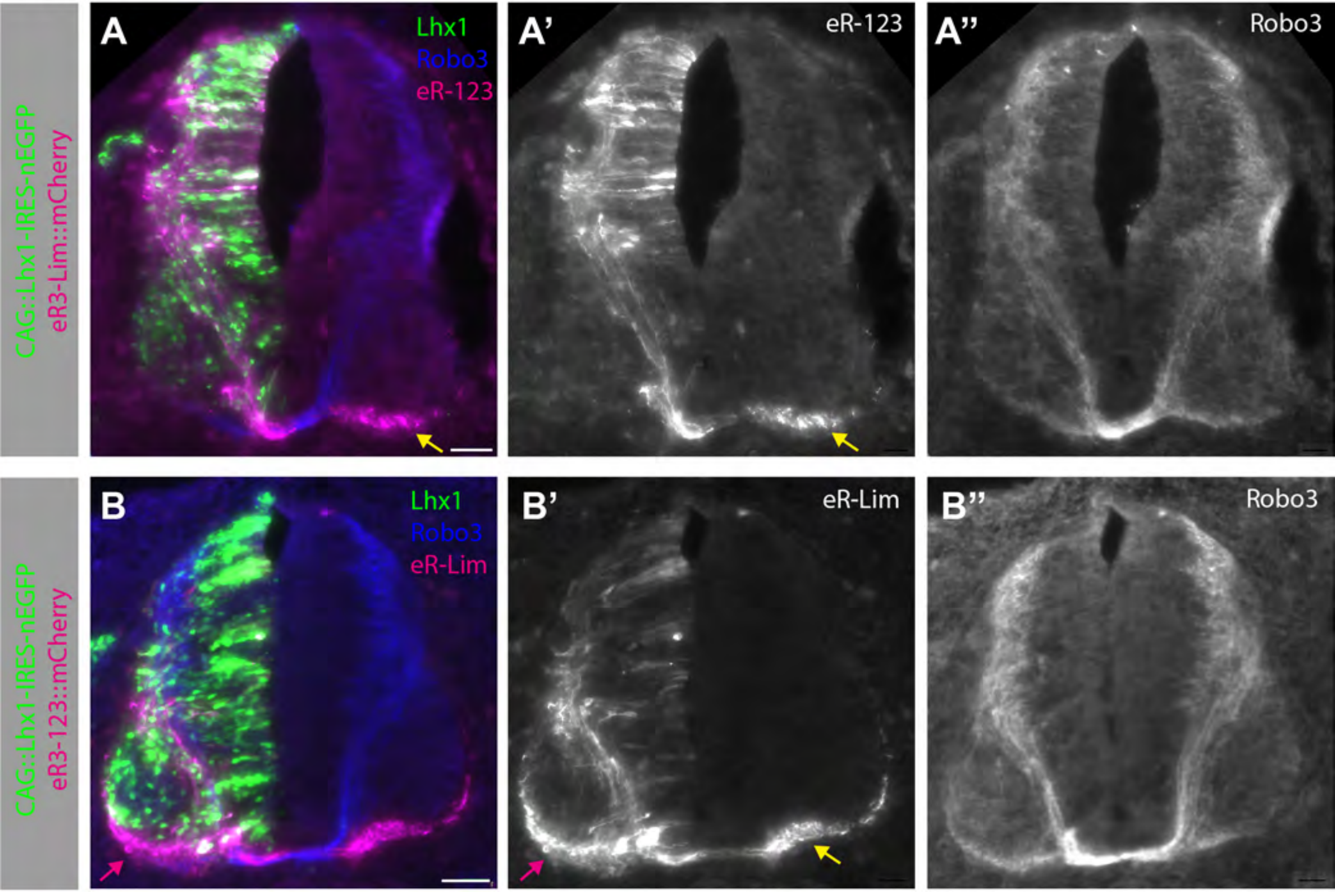
CAG::Lhx1-IRES-nEGFP
eR3-Lim::mCherry
A
Lhx1
Robo3
eR-123
A'
eR-123
A''
Robo3
CAG::Lhx1-IRES-nEGFP
eR3-123::mCherry
B
Lhx1
Robo3
eR-Lim
B'
eR-Lim
B''
Robo3
